# Slower cardiac-coupled cortical dynamics link depressive symptoms to inflexible affective updating

**DOI:** 10.64898/2026.08.11.744232

**Authors:** Jinwoo Lee, Kyungjin Oh, Junhyung Kim, Jiook Cha

**Affiliations:** Department of Psychology, Seoul National University, South Korea; Department of Psychology, University of California, San Diego, United States; Department of Psychiatry, Kangbuk Samsung Hospital, Sungkyunkwan University, South Korea; Department of Brain and Cognitive Sciences, Seoul National University, South Korea; Interdisciplinary Program of Artificial Intelligence, Seoul National University, South Korea; Institute of Psychological Science, Seoul National University, South Korea

## Abstract

Depression is marked by blunted affective responses to context, which interoceptive accounts trace to altered neural representations of bodily states. Yet this evidence mainly concerns response magnitude, not how quickly affect is updated when contexts change. Here we tested whether depressive symptom severity is related to delayed affective updating, and whether cortical dynamics tracking cardiac states account for this delay. To this end, we applied a movie-watching paradigm with independently defined contextual shifts, continuous affect ratings, electroencephalography, and electrocardiography in individuals spanning a continuum of depressive symptoms. Combining deep representation learning and a dynamical systems framework, we quantified how quickly (speed) and how sharply (angle) cardiac-coupled cortical representations reorganized at each shift. Greater symptom severity predicted longer latency to enter the context-congruent affective state across contextual shifts, regardless of valence. In a cross-sectional mediation analysis, slower speed, but not angle, accounted for this association. This mediation was specific to depressive symptoms, contextual shifts, and cardiac-coupled neural dynamics. These findings extend the embodied account of depression from blunted affective intensity toward its inflexible updating at moments of contextual shift, and offer a broadly applicable framework for quantifying brain-body dynamics across affective dysfunctions.

## Introduction

Affective flexibility, the capacity to update one’s affective state in alignment with shifting environments, is essential for navigating an uncertain world. Affect functions as a predictive signal that manages the ‘body budget’, enabling efficient actions in anticipation of upcoming events to maintain homeostasis^1–3^. Updating one’s affective state to changing contexts in a context-congruent manner is therefore indispensable for adaptive behavior, and is associated with adaptive outcomes such as resilience^4^ and subjective well-being^5^.

Mounting evidence suggests that depressive symptoms, across a continuous spectrum from subclinical to clinical severity, systematically degrade affective flexibility. Nevertheless, when, in what temporal pattern, and why such impairment arises remain open questions. Ecological momentary assessment studies have captured day-to-day affective persistence in depressive symptom severity by reporting elevated inertia of negative affect^6–8^. However, a large-scale meta-analysis recently demonstrated that such context-agnostic statistics offer little predictive power beyond mean affect intensity for psychological conditions^9^, motivating a shift toward ‘contextualized’ affect dynamics anchored to critical perturbation moments at which the affective state must be updated^10–12^. Additionally, whether depressive symptoms are linked to more rigid updating selectively toward positive contexts or generally across valences also remains debated^13–15^. Finally, the neurocomputational basis engaging in such (in)flexible updating of affective states has also not been empirically examined. A comprehensive understanding of how depressive symptom severity alters temporal architecture of affective updating therefore requires (1) testing whether depressive symptoms are related to delayed context-congruent affect updating under controlled contextual shifts, (2) examining whether this delay depends on the valence of the shifting context, and (3) identifying its neurocomputational substrate.

Addressing these questions calls for a theoretical framework that links depression-related inflexibility in affective updating to its neural basis. The interoception-centered perspective, which conceptualizes affect as the neural representation of bodily states, offers such a framework^16–18^. Within this view, an affective state arises as a posterior inference integrating the brain’s prior beliefs about bodily condition with likelihood of visceral signals. Thus, updating the affective state in response to a contextual change is characterized as a dynamic shift in neural representation of bodily states. Depression has been understood as a condition in which noisy visceral perception and over-weighted prior beliefs render this posterior update rigid^19–22^. The framework thereby predicts that the delayed affective updating associated with depressive symptoms should be accompanied by altered neural dynamics of bodily states during contextual shifts.

Testing this prediction requires an analytical approach capable of directly quantifying the temporal dynamics of body-related neural representations. For example, brain-heart interplay has been studied through heartbeat-evoked potentials that capture cortical responses to cardiac afferents^23,24^, or Bayesian generative modeling of interoceptive precision^25^. However, neither directly addresses how the neural representations of cardiac states evolve over time, particularly at the moment of contextual shifts. The dynamical systems framework has been proposed to quantify brain-body joint dynamics with trajectories in a latent neural space^26,27^, with nonlinear dimensionality reduction techniques proposed as a useful approach for their estimation^28,29^. Within this view, affective updating is fundamentally a movement from one region of the cardiac-coupled neural space to another, and such movement can be quantified by two first-order geometric features of the trajectory: its speed and angle. These two features have recently been validated as representative measures showing dissociable dynamical patterns across cognitive processes^30,31^. Therefore, we exploratorily investigated the roles of two dynamic motifs: whether inflexible affective updating linked to depressive symptoms is selectively related to the slow speed or small angle of neural dynamics, or both.

To operationalize these predictions, we combined continuous affect ratings with electrophysiological recordings during movie watching, a paradigm that affords three converging strengths. First, narrative arcs reliably elicit a sequence of affective contexts within a single session^32,33^, which enables a within-subject comparison of affective updating dynamics across valences and thus a critical test of whether depression-related inflexibility is valence-specific or valence-general. Second, because all participants view the same movie, individual differences in affective updating can be compared under a shared and contextualized environment^34,35^. Since contextual shift points can be defined objectively through independent annotators, it allows one to quantify the deviation of one’s affective trajectory from the normative one while avoiding the self-referential context definition inherent to ecological momentary assessment. Third, continuous affect ratings, electrophysiological signals, and film timestamps share a timeline, allowing affective updating and neural dynamics aligned with cardiac states to be resolved at second-to-second resolution, the timescale at which depression-related alterations in affect dynamics are most effectively measured^36,37^.

Our study performs the following confirmatory hypothesis tests: (1) context-congruent affective updating is delayed as a function of depressive symptom severity, and (2) this association is statistically mediated by blunted dynamics of the cardiac-coupled cortical representation at contextual shifts. Additionally, our study includes the following exploratory analysis: (3) Does the relationship between depressive symptoms and affective updating depend on the valence of the shifting context? and (4) How do the speed and angle of cardiac-coupled cortical dynamics account for the association between depressive symptoms and delayed affective updating?

To investigate these, we combined film watching with concurrent continuous affect ratings, electroencephalogram (EEG), and electrocardiogram (ECG). We applied a dynamical systems framework to the EEG-ECG signals to extract each individual’s trajectory speed and angle for every timepoint and tested whether these features statistically mediate the association between depressive symptoms and latency of affective transition at the shifts of movie contexts. By directly measuring the second-to-second dynamics of contextual shift moments, this study extends the embodied account of depressive symptoms from blunted magnitude toward inflexible affect dynamics.

## Results

### The movie provides rich contexts associated with diverse subjective affect

Forty-three young adults spanning a continuous spectrum of depressive and anxiety symptoms completed the experimental protocol (**Figs 1a-b** and **Methods**); BDI-II total scores ranged from 0 to 29, with 21% of participants exceeding the clinical cutoffs. After providing demographic information and psychological assessments, participants underwent a resting-state measurement with 19-channel EEG and a single-channel ECG. To familiarize them with real-time affect rating using Evaluative Space Grid^38^, participants practiced reporting their on-going subjective feelings while watching a short trailer of the movie *Up*. For the main task, we used *One Small Step*^39^, an Academy Award-nominated 7-min animation validated to efficiently evoke rich affective feelings over time^40^. During the movie, we recorded continuous ratings of positive and negative affect along with synchronized EEG/ECG signals, followed by a post-trial questionnaire.

**Fig 1.**
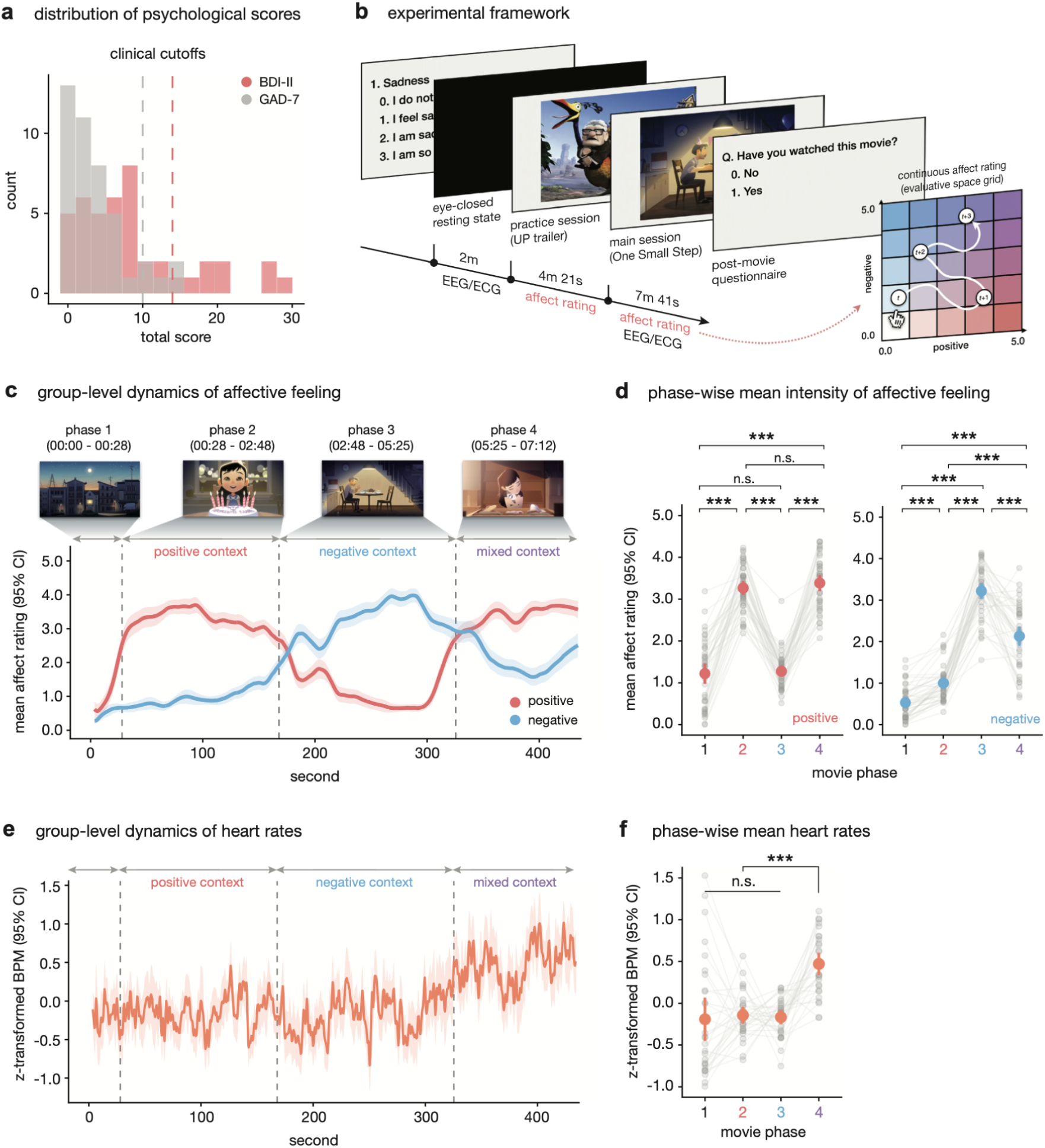
Movie-watching paradigm and affective characterization of movie contexts. **a,** Distribution of two psychological measures. Dotted lines represent the clinical cutoff scores for each questionnaire (i.e., 14 for BDI-II and 10 for GAD-7). **b,** Experimental framework. **c and e,** Group-level dynamics of affective ratings and heart rate. Each graph shows the group-level mean of self-reported affect and z-scored heart rate (beats per minute) in the main experiment (*N* = 43). The gray dotted line denotes the time point of contextual changes annotated by independent raters (*N* = 18). **d and f,** Phase-wise mean intensity of affective ratings and heart rate. Each gray point represents an individual data point. Statistical differences were assessed using paired two-sided *t*-tests. \**P*_FDR_ < .05; \*\**P*_FDR_ < .01; \*\*\**P*_FDR_ < .001.

To identify contextual transition points of the movie without circularity, we recruited independent annotators who reported real-time affect using the same protocol (*N* = 18; **Supplementary Fig 1**). We identified four movie phases with three context transitions corresponding to key narrative and affective shifts: the introduction of the protagonist girl’s family (phase 1), the pursuit of her dreams (phase 2), the loss of a main caregiver (phase 3), and ultimate achievement following grief (phase 4). Based on these transition timepoints, we characterized resultant four movie phases as neutral, positive, negative, and mixed contexts.

The average affect ratings from the main experimental group closely tracked these identified transition points (**Fig 1c**). Within individuals, positive affect intensity was significantly higher during positive and mixed phases than neutral and negative ones, while negative affect intensity followed a descending order from negative, mixed, positive, to neutral phases (**Fig 1d** and **Supplementary Table 1** for statistics). Beyond self-reported affect, group-level heart rate (beats per minute, BPM) significantly increased during the last phase, the affective climax where grief and joy coexist, compared to preceding phases (**Figs 1e-f** and **Supplementary Table 1** for statistics).

To address potential concerns regarding cognitive load and the limited fidelity of continuous affect ratings, we compared our participants’ group-level affect dynamics with previous findings using the same movie stimulus (the ref.^40^; *N* = 27). Despite differences in participants’ cultural backgrounds (East Asian vs. North American), we found high temporal correlations across all affect dimensions: positive: *r*(429) = .900, 95% CI = [.881, .917], *P*_FDR_ < 2.2×10^-16^; negative: *r*(429) = .928, 95% CI = [.914, .940], *P*_FDR_ < 2.2×10^-16^; mixed: *r*(429) = .578, 95% CI = [.511, .637], *P*_FDR_ < 2.2×10^-16^ (**Supplementary Fig 2**).

These results demonstrate that the movie stimulus efficiently evokes diverse affective contexts in a naturalistic setting, supporting the suitability of our paradigm for capturing individual differences in real-time affective updating across changing contexts.

### Greater depressive symptom severity predicts consistent delays of affective updating across contextual shifts

We next examined whether higher depressive symptom severity is associated with delay in the update of affective state in response to changes in movie contexts. To clearly define ‘the update of affective state’, we first categorized each participant’s affect ratings at every timepoint into one of four states - neutral, positive, negative, or mixed - using a threshold of 2.5, the midpoint of the Evaluative Space Grid. Based on each individual’s affective state sequence, we calculated the transition latency, defined as the time taken from a contextual transition to the first entry into the affective state congruent with the corresponding context (**Fig 2a** and **Methods**). For example, for transitions into Phases 2, we measured the latency to reach positive states. Across three transitions, participant’s latency showed significant and fair reliability (ICC(3, 1) = .287, 95% CI = [.100, .490], *F*(42, 84) = 2.21, *P* = .001; 43 participants × 3 phases = total 129 observations), which remained robust regardless of the threshold applied for discretizing affective states (**Fig 2b** and **Supplementary Table 2** for statistics). Also, we confirmed that, with the exception of a single case in which one participant failed to enter the negative state upon transitioning to Phase 3, all participants successfully entered the phase-congruent state in every phase.

**Fig 2.**
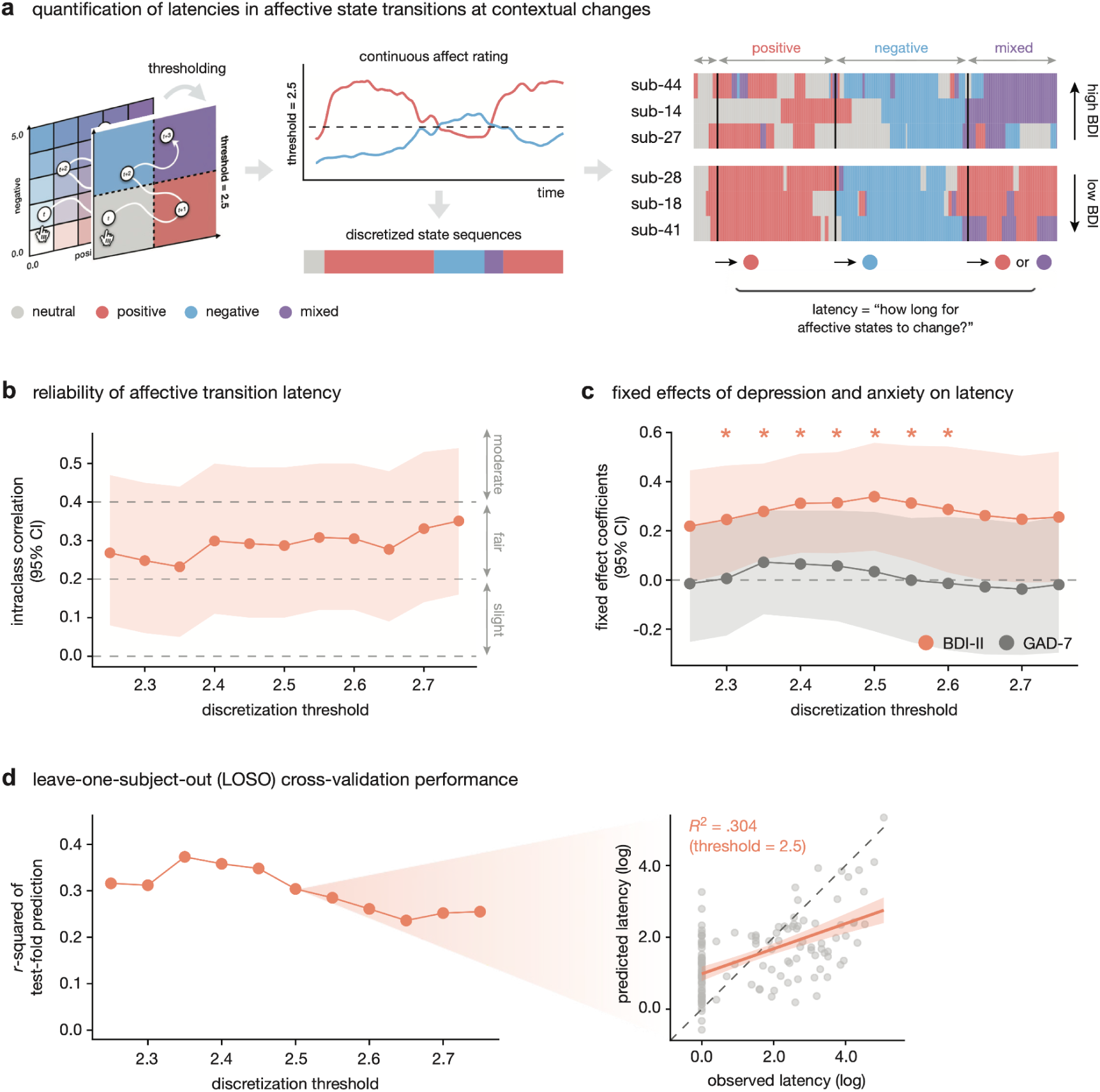
A latency of affective state transition at the contextual changes and its association with depressive states. **a,** Quantification of latencies in affective state transitions at contextual changes. By applying a threshold of 2.5 to the evaluative space grid, we discretized each participant’s affective dynamics into state sequences. Latency was defined as the time elapsed from each contextual change point (i.e., black solid line) to the first entry into the affective state dominant within that phase. **b,** Reliability of affective transition latency. The reliability of the latency calculated from each of the three transitions was assessed using the intraclass correlation coefficient (ICC). The interpretation of the ICC followed the conventional criteria of the ref.^101^. The shaded area represents a 95% confidence interval. ICCs were iteratively estimated for discretization threshold parameters ranging from 2.25 to 2.75 in 0.05 increments. **c,** Fixed effects of depression and anxiety on latency. Shaded areas indicate 95% confidence intervals for the fixed effects of each psychological measure in the linear mixed-effects models. Fixed effect coefficients were iteratively estimated for discretization threshold parameters ranging from 2.25 to 2.75 in 0.05 increments. Orange asterisks indicate significant fixed-effect estimates with thresholds ranging from 2.3 to 2.6 at *α* = .05. **d,** leave-one-subject-out cross-validation performance. On the left, the *R*-squared in the held-out folds across discretization thresholds were visualized. On the right, a scatterplot of the predicted versus observed log-transformed latency at a threshold of 2.5 is presented. The diagonal dashed line indicates perfect prediction.

We tested if BDI-II scores explain latency of affective transition after controlling for sex, age, phase type, and per-phase affective peak intensity using a linear mixed-effects model with random intercepts. Since a weaker affective response crosses a fixed threshold later, we included per-phase affective peak intensity as a covariate, so that transition latency clearly indexes the timing rather than the magnitude of the affective response. Also, to account for the skewed distribution of latency, we used the log-transformed value of the latency as an outcome variable (see **Methods**).

Higher BDI-II scores were significantly associated with the longer latency (fixed-effect *β* = .339, 95% CI = [.119, .558], *P* = .003, BF_10_ = 5.717), and this relationship remained robust across the discretization thresholds of affective states from 2.30 to 2.60 (**Fig 2c** and **Supplementary Table 3**). In contrast, models using GAD-7 as a fixed regressor failed to explain transition latency, which was also robust to the choice of threshold (fixed-effect *β* = .034, 95% CI = [−.209, .277], *P* = .783, BF_10_ = .091; **Fig 2c** and **Supplementary Table 3**). Diagnostic checks for the BDI-II model confirmed that key assumptions for residual normality, homoscedasticity, linearity, and random intercept normality were all met (**Supplementary Fig 3**).

To assess predictive validity of the BDI-II model, we conducted leave-one-subject-out (LOSO) cross-validation (see **Methods**). The model predicted latencies for held-out participants across all transitions (test *R*^2^_predicted~observed_ = .304; test *r*_predicted~observed_ = .556, *P* = 7.6×10^-12^), and this performance was robust across discretization thresholds (**Fig 2d**).

Last, we exploratorily examined whether the relationship between depressive symptom severity and affective transition latency was modulated by the valence of each context shift or not. To investigate this, we compared the above main effects model against a new model adding an interaction term between BDI-II score and phase type (see **Methods**). Model comparison provided moderate evidence in favor of the main effect model without the interaction term (BF_10_ = .126) across all discretization thresholds (**Supplementary Fig 4**), which indicates that the relationship between depressive symptoms and delayed affective updating was consistent across valence shifts in contextual changes.

These findings indicate that depressive symptoms predict delays in affective updating to changing contexts in a valence-general manner.

### Deep representation learning reliably models cardiac-coupled neural representations in a biologically plausible manner

To model latent neural representations coupled to cardiac states during movie watching, we first identified a subset of participants with high-quality EEG and ECG data (*N* = 30). We tested whether inclusion (or exclusion) in this analysis had a confounding effect on the BDI-II score and the latency of affective transitions by comparing these variables between the included and excluded groups; no statistically significant differences were found for any variable (all *P*s ≥ .382 and the absolute value of rank-biserial *r*s ≤ .172; **Supplementary Fig 5**).

Here, we applied CEBRA^41^, which performs nonlinear dimensionality reduction of neural signals by leveraging auxiliary variables. Such supervised approach makes it a powerful tool for isolating neural representations of specific affective or autonomic-cortical axes from complex sensory inputs^28,29,42^. We fitted a multi-participant model using 90 EEG features (i.e., 18 channels × 5 band frequencies) as inputs and temporally aligned beats-per-minute (BPM) time-series as the auxiliary variable to encode cardiac states onto a shared neural space (**Fig 3a** and **Methods**). A 9-dimensional embedding space was selected for main analyses based on its highest contrastive performance of BPM values among all tested dimensionalities ranging from 2 to 9 (**Fig 3b**).

**Fig 3.**
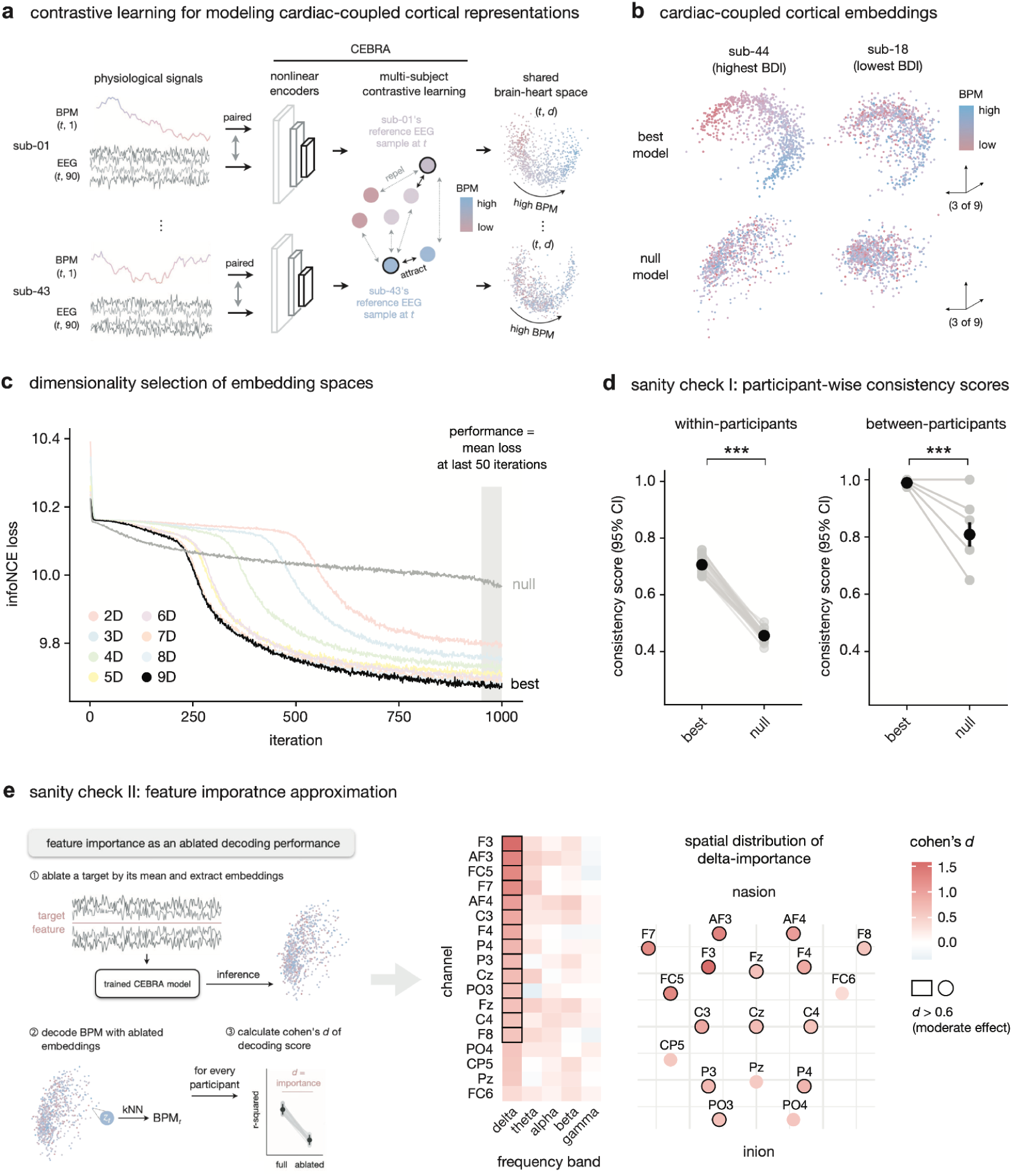
Framework of modeling cardiac-coupled cortical representation and sanity checks. **a,** Contrastive learning for modeling cardiac-coupled cortical representations. We applied CEBRA-Behavior to each participant’s EEG and ECG time series. Using multi-subject training, all participants’ resulting cardiac-coupled cortical space share common axes and representational geometry. **b,** Cardiac-coupled cortical embeddings. Embeddings from two representative participants are shown. Unlike the null model, the best-performing nine-dimensional model encodes beats per minute (BPM) along a gradient. **c,** Dimensionality selection of embedding spaces. To determine the optimal dimensionality of the embedding spaces, we compared contrastive performance across CEBRA models ranging from two to nine dimensions, based on the mean InfoNCE loss over the last 50 iterations. **d,** Sanity check I: participant-wise consistency score. Each gray point represents a participant’s consistency score. Statistical differences between the best and null models were tested using paired two-sided t-tests. \**P* < .05; \*\**P* < .01; \*\*\**P* < .001. **e,** Sanity check II: feature importance approximation. We assessed the importance of a total of 90 features in the CEBRA training process using ablation-based decoding analyses. Feature importance was quantified by the extent to which decoding performance for BPM degraded when each feature was ablated (i.e., averaged across all time points for each participant) and then fed into the best model.

We validated the reliability and biological plausibility of our modeling approach. First, we assessed the consistency of the embedding geometry. Consistency quantified the structural similarity between models (within-participants) and participants (between-participants) by measuring how accurately an embedding from one model (or participant) could linearly predict another (see the ref.^41^ and **Methods**). The actual models captured significantly more consistent representation geometries than null models trained on shuffled BPM, both within-individuals (within-participants consistency: *t*(29) = 37.339, 95% CI = [.24, .26], *P* < 2.2×10^-16^) and across-individuals (between-participants consistency: *t*(29) = 9.267, 95% CI = [.14, .22], *P* < 2.2×10^-16^; **Fig 3d**).

Second, we evaluated whether the model relied on neurobiologically relevant features to construct the cardiac-coupled neural space. We quantified the importance of each EEG feature for constructing the latent neural space based on how much its ablation degraded the decoding performance of embeddings for BPM (**Fig 3e** and **Methods**). We found that low-frequency delta bands (1-4 Hz), particularly in front-central channels (e.g., F3, AF4, FC5), exhibited the highest importance (**Fig 3e**). This pattern aligns with prior findings highlighting front-central delta power as a causal signature of top-down control over cardiac signals^43^ and as an active modulator in autonomic regulations^44^.

Collectively, these findings demonstrate that our model not only captures geometry of cardiac-coupled neural representations reliably, but also leverages neural features related to encoding of cardiac states, providing a neurophysiological foundation for analyzing cardiac-coupled cortical dynamics.

### Slower cardiac-coupled cortical dynamics statistically mediate the association between depressive symptoms and delayed affective updating

Using the 9-dimensional embedding space, we quantified the geometric dynamics of cardiac-coupled neural trajectories during contextual shifts with speed and angle. We then tested whether these dynamic motifs statistically mediate the association between depressive symptoms and the delays in affective transitions. For each contextual transition, we defined a ‘transition window’ with a window size parameter *k* (seconds; i.e., [*-k, +k*] from the transition timepoint) and calculated the average embedding speed and angle across all timepoints in three transition windows (**Fig 4a** and **Methods**). Here, speed represents the distance the embedding moved per unit time, quantifying the magnitude of the brain-heart embedding changes during the contextual transition. Angle reflects how sharply the embedding turned from its previous trajectory in the embedding space, representing the directionality or qualitative difference of the brain-heart embedding change during the contextual transition. For both metrics, the group in the top 10% of BDI-II scores showed more attenuated dynamics throughout the movie than the group in the bottom 10% (**Fig 4b**).

**Fig 4.**
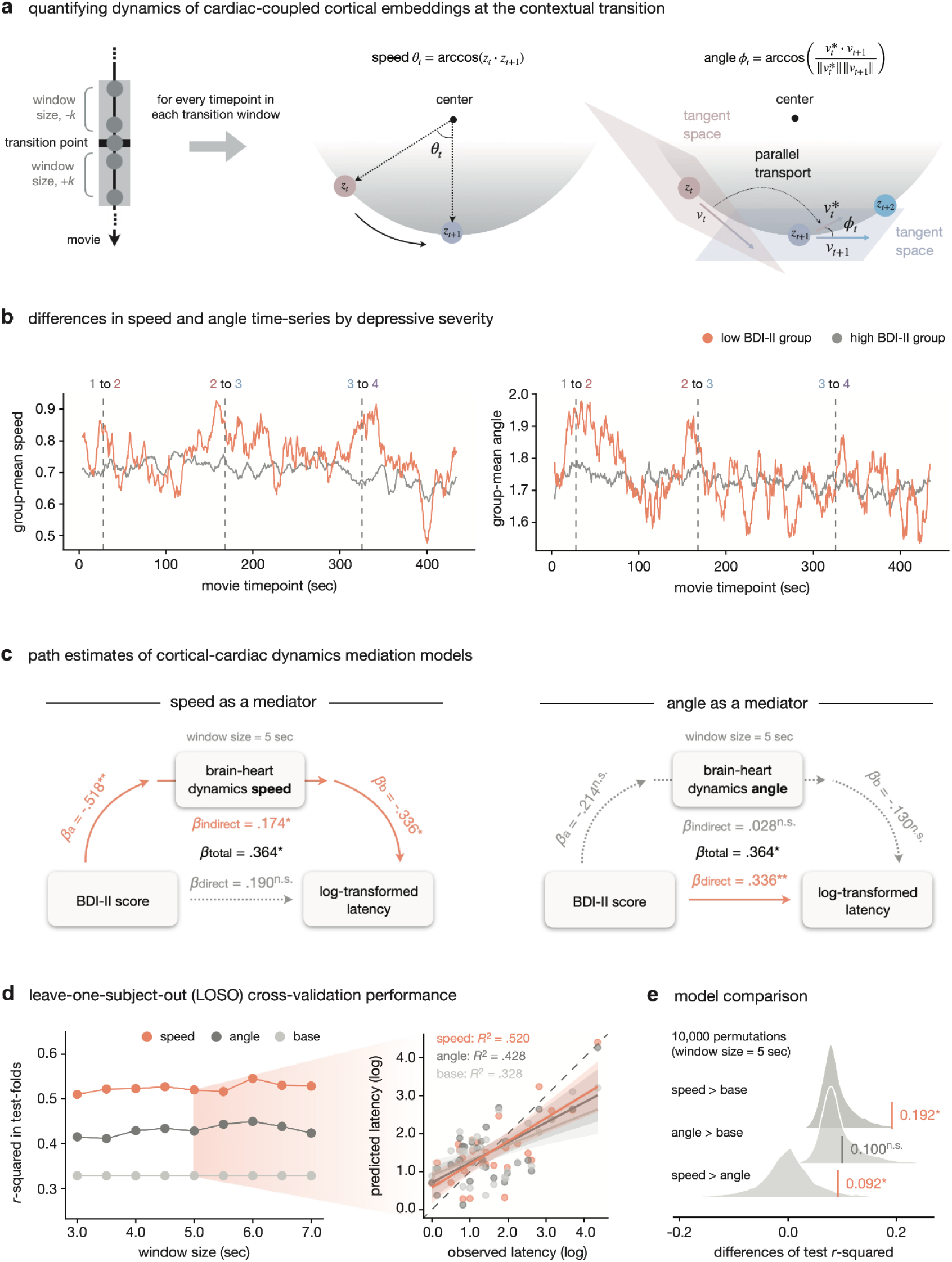
Mediation of brain-heart dynamics linking depressive severity and affective updating. **a,** Quantifying dynamics of cardiac-coupled cortical embeddings at contextual transition. For every timepoint, we calculated the speed and angle of brain-heart embeddings from the best model and averaged each feature across all timepoints within three transition windows. We used the resulting speed and angle values as representative brain-heart dynamics features. **b,** Differences in speed and angle time series by depressive severity. The left and right panels show the mean speed and angle time series, respectively, for the top (gray) and bottom (orange) 10% of participants based on BDI-II total scores across the entire movie. Vertical dotted lines indicate transitions in affective context. **c,** Path estimates of cortical-cardiac dynamics mediation models. We tested the mediation model which used a speed and angle as a mediator, respectively, while controlling for sex, age, and phase-averaged peak intensity of context-congruent affect. All statistics were calculated with a window size of 5s with 10,000 bootstrap. \**P* < .05; \*\**P* < .01; \*\*\**P* < .001. **d,** Leave-one-subject-out (LOSO) cross-validation performance. On the left, the test *R*-squared for each window size parameter are shown as a performance metric: speed- (orange), angle-mediator (dark gray), and baseline models (light gray). On the right, a scatterplot of the predicted versus observed log-transformed latency at a window size of 5s is presented for each model. The diagonal dashed line indicates perfect prediction. **e,** Model comparison. Based on the preceding LOSO cross-validation results, we tested the hypothesized ranking of predictive performance, speed-mediator > angle-mediator > and base models, using 10,000 permutations for each pairwise comparison (30,000 in total). The gray distribution in each pairwise test represents the null distribution of the difference in *R*-squared, and the vertical line indicates the observed difference. Orange lines show significant differences at *α* = .05.

Across the three context transitions, the speed and angle of each participant demonstrated significant and moderate reliability within a *k* = 5 window size (speed: ICC(3, 1) = .323, 95% CI = [.10, .56], *F*(29, 58) = 2.432, *P* = .002; angle: ICC(3, 1) = .402, 95% CI = [.18, .62], *F*(29, 58) = 3.015, *P* < .001; 30 participants × 3 phases = total 90 observations for each metric), and this reliability was robust to variations in parameter *k* (**Supplementary Table 4**).

We performed cross-sectional mediation analyses with *z*-scored BDI-II score as the input variable, *z*-scored speed or angle as the mediator, and log-transformed mean affective transition latency as the outcome, while controlling for age, sex, and phase-averaged peak intensity of context-congruent affect (see **Methods**). Within a *k* = 5 transition window, embedding speed significantly mediated the association between BDI-II scores and affective transition latency (*β*_indirect_ = .174, 95% CI = [.036, .416], *P* = .041; **Fig 4c**). This mediation was robust across different window size parameters ranging from 3.0 to 7.0 seconds (**Supplementary Table 5**). In contrast, the embedding angle did not show significant mediation across the window size parameters (under *k* = 5 window: *β*_indirect_ = .028, 95% CI = [−.022, .190], *P* = .542 from 10,000 bootstrap; **Fig 4c** and **Supplementary Table 5**).

Finally, to evaluate whether the speed-mediator model could predict held-out participants’ latency, we conducted LOSO cross-validation (see **Methods**). Regardless of the window size parameters, predictive performance was consistently highest for the speed-mediator model, followed by the angle-mediator model and the base model, which included only covariates (under *k* = 5 window: *R*^2^_speed_ = .520, *R*^2^_angle_ = .428, *R*^2^_base_ = .328; **Fig 4d**). Under the *k* = 5 window condition, the speed-mediator model showed significantly greater performance than the other two models (*ΔR*^2^_speed-base_ = .192, *P* = .023; *ΔR*^2^_speed-angle_ = .092, *P* = .041; from 10,000 permutations; see **Fig 4e**), whereas the angle-mediator and base models did not differ significantly (*ΔR*^2^_angle-base_ = .100, *P* = .298; from 10,000 permutations; **Fig 4e**).

These findings consistently indicate that slower speed of cardiac-coupled cortical dynamics at contextual transitions accounts for the association between depressive symptoms and delayed affective updating, whereas trajectory angle does not.

### The mediation is specific to depressive symptoms, contextual transition, and cardiac-coupled neural dynamics

To evaluate the specificity of the observed mediation, we conducted three sets of control analyses focusing on input, modality, and timing dimensions (**Fig 5a**). First, we replaced the input variable with GAD-7 scores to test whether the mediation effect is specific to depressive symptom severity rather than anxiety severity. Second, we tested whether cardiac-coupled neural dynamics are uniquely responsible for this mediation by training a ‘CEBRA-time’ model which uses timepoints indices as an auxiliary variable instead of BPM to use pure neural dynamics as a mediator. Last, to verify whether the effect is specific to the moments of contextual transitions, we calculated embedding dynamics within ‘control windows’ centered at the midpoint of each movie phase that did not include contextual shifts.

**Fig 5.**
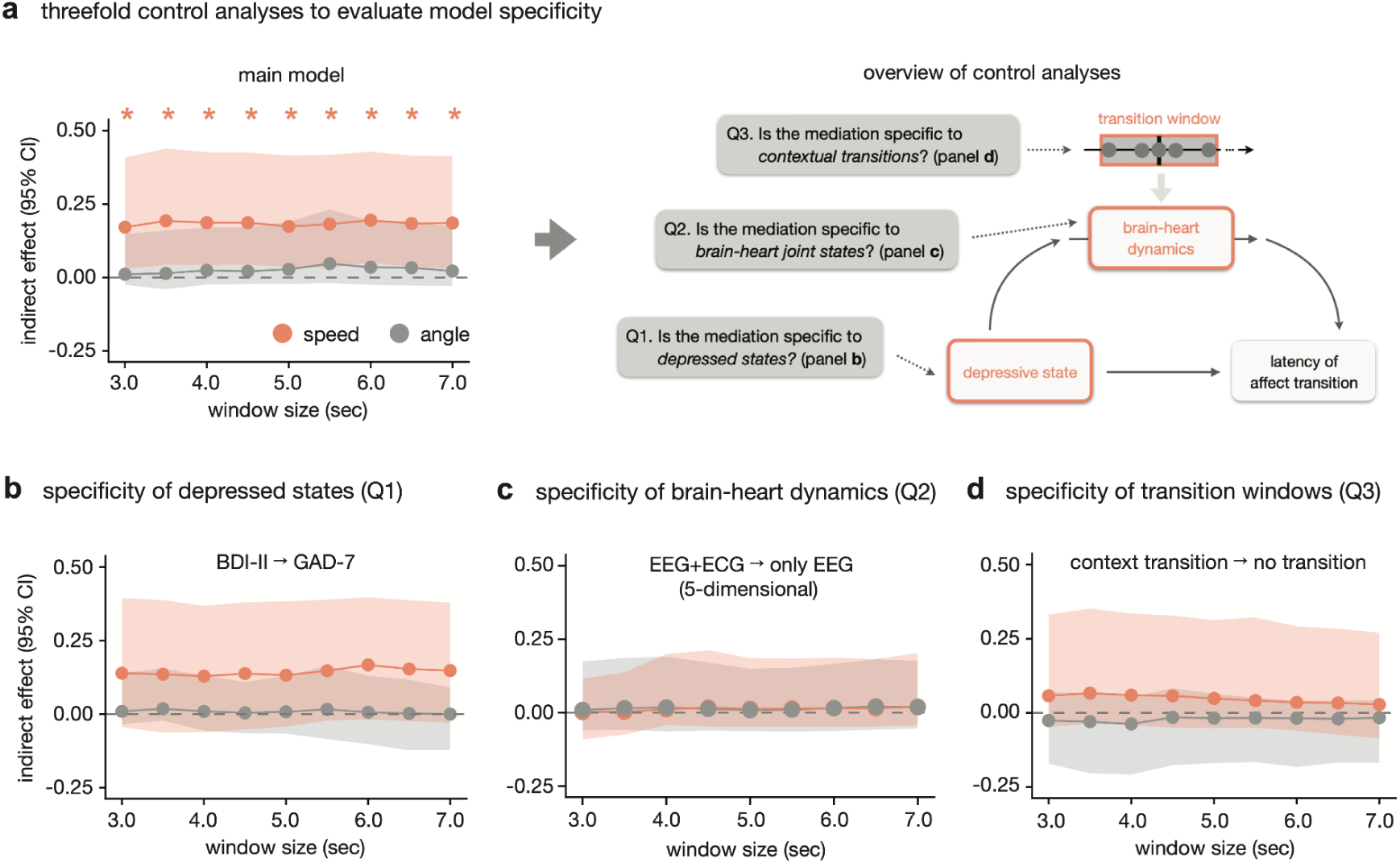
Threefold control analyses to evaluate specificity of mediation model. **a,** Threefold control analyses to evaluate model specificity. On the left, beta coefficients of indirect effect in the main model and their 95% confidence intervals are visualized across window size parameters for each mediator. Asterisks indicate significant estimates at *α* = .05. On the right, a conceptual overview of our control analyses to the main results was shown. **b-d,** Specificity checks for depressed states (input), brain-heart joint states (modality of mediator), and contextual transitions (timepoint of mediator). All confidence intervals of indirect effects were estimated using 10,000 times bootstrapping for window sizes ranging from 3 to 7s in 0.5s increments.

First, in the mediation model using GAD-7 total score as the input variable, GAD-7 score did not show significant indirect effects through either embedding speed or angle (speed under *k* = 5 window: *β*_indirect_ = .132, 95% CI = [−.043, .383], *P* = .248; angle under *k* = 5 window: *β*_indirect_ = .008, 95% CI = [−.066, .130], *P* = .898). This lack of significant mediation was consistent across all tested window sizes (**Fig 5b** and **Supplementary Tables 6**).

Second, we trained a CEBRA-time model to assess whether pure neural dynamics, rather than cardiac-coupled dynamics, could account for the mediation. A 5-dimensional model was selected as the best embedding space model (**Supplementary Fig 6**). Both speed and angle derived from these embeddings did not significantly mediate the link between BDI-II score and affective latency (speed under *k* = 5 window: *β*_indirect_ = .013, 95% CI = [−.048, .186], *P* = .757; angle under *k* = 5 window: *β*_indirect_ = .008, 95% CI = [−.063, .149], *P* = .803). These findings remained consistent across all window sizes (**Fig 5c** and **Supplementary Table 7**). When the embedding dimensionality was set to 9-dimensions to match the dimensionality used in the main cardiac-coupled analysis, the mediation was still not significant regardless of the window size parameter (**Supplementary Fig 7**).

Third, when using embedding dynamics from control windows centered at phase midpoints as mediators, neither speed nor angle significantly mediated the relationship between BDI-II score and affective latency (speed under *k* = 5 window: *β*_indirect_ = .048, 95% CI = [−.052, .313], *P* = .468; angle under *k* = 5 window: *β*_indirect_ = -.018, 95% CI = [−.170, .065], *P* = .587). This lack of significance was robust to all tested window sizes (**Fig 5d** and **Supplementary Tables 8**).

Taken together, these three control analyses consistently indicate that the mediation is specific to depressive symptom severity as the predictor, the cardiac-coupled cortical dynamics as the mediator, and the moments of contextual transitions.

## Discussion

In this study, we investigated whether and how depressive symptom severity is associated with delayed affective updating during contextual shifts, and explored its neurocomputational substrate. We combined the interoception account with a dynamical systems framework to model cardiac-coupled cortical dynamics. We applied deep representation learning to derive a cardiac-coupled cortical representation for each participant, and quantified its dynamics using two trajectory features; speed and angle. We then tested whether these metrics statistically mediated the association between depressive symptoms and affective transition latency. We observed that greater depressive symptom severity predicted longer affective transition latency across all three contextual shifts regardless of shift valence, indicating the valence-general nature of such inflexibility. This association was statistically mediated by slower speed of cardiac-coupled cortical dynamics; an analogous test of dynamics angle did not reach significance. This pattern suggests that more severe depressive symptoms are related to a slower rate of affective updating, accompanied by slower cortical dynamics. Three control analyses indicated that this mediation was specific to depressive symptoms (not anxiety symptoms), to contextual transition (not non-shift moments), and to cardiac-coupled cortical dynamics (not pure neural dynamics). Together, these findings provide insights into the embodied account of depression-related affective inflexibility, promoting a contextualized and dynamics-based understanding of how depressive symptoms interact with inflexible affect dynamics.

Our behavioral finding of valence-general inflexibility in affective updating extends the Emotion Context Insensitivity framework into a previously underexplored temporal dimension. Traditional emotion context insensitivity studies have demonstrated that depressive symptom severity is associated with reduced magnitude of response to both positive and negative emotional stimuli in self-reported^45^ and neurophysiological^46^ measures. Motivational withdrawal, which frames depression as a chronic resource conservation strategy that dampens approach and avoidance motivation, has been proposed as one of the prominent cognitive mechanisms^47–49^. Building on this account, we propose that the motivational disengagement related to depressive symptoms may impair not only the intensity of affective reactivity, but also the temporal dynamics of affective updating during contextual shifts. When combined with measurements of motivational engagement (e.g., reward positivity, pupil dilation) and direct examination of their relationship with latency of affective updating, our approach could reveal a temporal profile of context insensitivity that has been largely unexplored and its underpinning cognitive basis.

Beyond this temporal characterization at the behavioral level, our neural findings provide initial insight into the cortical dynamics accompanying such inflexibility. We interpret the BPM-supervised latent EEG embedding as a cardiac-coupled cortical representation, which has been conceptualized as one critical substrate for valenced affective experience across multiple emotion theories^2,16,18^. Supporting this interpretation, our feature attribution analysis identified low-frequency power in frontocentral channels as key features for the cardiac-coupled representations. Frontocentral delta oscillations have been previously linked to top-down predictions of bodily signals^1,50,51^. Moreover, recent neuromodulation work provides causal evidence that frontal delta synchrony modulates heartbeat perception^43^. We acknowledge that residual cardiac artifacts, which may persist even after removal of heartbeat-related components, could drive delta-band importance. However, the findings that the QRS complex or pulse contamination is broadband^52,53^ and left-lateralized^54,55^ argue against an account in which the importance of bilateral delta oscillations merely reflects residual cardiac contamination.

Building on this representation, we then asked how its dynamics reorganize at contextual shifts. Within the dynamical systems framework, embedding speed quantifies how rapidly this cortical representation changes at moments of contextual change. Our finding that speed, but not angle, mediated the association resonates with conceptual proposals that characterize depression as involving ‘speed’ deficits across perception, affect, and motor control rather than deficits in their quality per se^56^. This speed-specific slowing is further consistent with computational accounts that conceptualize depressive symptoms as involving impaired precision-weighting of bodily signals^25,57–59^. Sluggish embedding speed can be understood as delayed posterior updating, in which precision impairments related to depressive symptoms may be linked to slower integration of new contextual evidence with bodily states into the cortical representation. Our trajectory-based approach thereby complements existing Bayesian framework for brain-heart interplay in depression: whereas Bayesian generative modeling estimates a static precision parameter (e.g., the ref.^25^), our framework characterizes the continuous evolution of cortical-cardiac dynamics as it unfolds across naturalistic contextual shifts.

In our trajectory-based framework, the dissociation between speed and angle offers a more nuanced perspective on how depressive symptoms relate to inflexible affective updating. Speed quantifies the magnitude of state movement per unit time, whereas angle captures the directional change between consecutive trajectory segments; the two are thus separable. Because speed, but not angle, significantly mediated the association, individuals with more severe depressive symptoms may retain information about the direction in which their affective states should be updated while moving more slowly toward it. Our behavioral data support this reading most directly: regardless of symptom severity, all participants eventually entered the phase-congruent affective state, with symptom severity linked to latency rather than distorted direction. Adjacent depression research offers indirect support. Studies combining the probabilistic reward task with drift diffusion modeling find that depressive symptom severity is associated with slower drift rates, but not with differences in directional bias toward particular choices^60–62^, which is analogous to our speed/angle dissociation. Given recent conceptualization of affective state entry as a decision-making process^63–65^, our findings yield a testable hypothesis: that persistent affect in depression, such as emotional inertia, reflects slower transition speed rather than altered reorientation.

Three control analyses delineated what drives and bounds the speed-based mediation, testing its specificity to the modality of the mediator, the timing of its operation, and the psychological state it indexes. We consider the modality and timing controls here, returning to the anxiety dissociation below. First, when the cardiac-coupled cortical dynamics were replaced by pure neural dynamics, the indirect effect was near-zero. This indicates that the mediation depends on cardiac information integrated into cortical representations rather than on generic neural dynamics, adding empirical support for embodied accounts of affect^16–18^. Because this control model was based on identical EEG signals without cardiac supervision, the result further argues against an account in which residual artifacts inherent in the EEG drive the observed pattern. Second, the mediation was attenuated when dynamics were computed outside the transition period. This supports a contextualized view of when affective inflexibility emerges^10–12,66^, an aspect less accessible to ecological momentary assessment approaches that rely on context-agnostic random sampling of timepoints. Our findings thus motivate refining the unit of analysis in affective chronometry from trait-like aggregates to context-locked states.

The last finding that replacing BDI-II scores with GAD-7 scores as the input left the indirect effect non-significant, pointing to a degree of specificity for depressive rather than anxious symptoms. We interpret this specificity cautiously: although the anxiety mediation did not reach significance, its point estimate was comparable in magnitude to that for depression, so the two cannot yet be firmly distinguished on the present data. That said, the direction of our results is at least conceptually consistent with computational accounts in which the two conditions differ in their interoceptive signatures. While anxiety is linked to heightened, volatile reactivity to bodily signals, depression with sluggish processing of them^22,67,68^. Under this view, this interpretation provides an open question: anxiety might be characterized by distinct dynamical features, such as instability rather than slowing.

More broadly, a dynamical perspective may help address long-standing inconsistencies in interoception research. Studies that relate internalizing symptoms to perceptual accuracy, a widely used static interoceptive measure, have yielded mixed results^69–71^. This motivates proposals that interoceptive processing can be conceptualized along multiple axes, including subjective confidence^72,73^ and rhythmic properties^51^. Our study contributes one such axis; the dynamic profile of cardiac-coupled cortical representations, which may help resolve heterogeneity within internalizing symptoms with further validation.

Several limitations of our study warrant explicit consideration. First, our transition latency measure assumes that all participants register the externally defined contextual shift at the same moment, but studies on event segmentation have reported that several psychiatric conditions are related to an altered perception of event transitions^74,75^. Although our control analyses show that brain-heart dynamics differ specifically around the annotator-defined transition windows, future work needs to test the difference in dynamical patterns between externally and internally defined contextual shifts. Second, our cross-sectional design cannot establish the causality of the observed mediation. Even where speed statistically mediated the association, we cannot infer that slowed cortical dynamics bring about delayed affective updating: reverse pathways, in which affective inflexibility shapes cortical dynamics, and unmeasured confounds (e.g., arousal or attention) remain possible. This limit also leaves the mediator’s interpretation underdetermined. The slower speed could equally reflect reduced precision-weighting of cardiac afferents or over-weighted prior. Adjudicating both the direction of the effect and computational basis behind it will require designs that go beyond statistical mediation, such as longitudinal measurement or pharmacological manipulation. Third, we only focused on cardiac signals as the bodily modality, but interoception encompasses multiple channels (e.g., respiratory, gastric) whose perceptual abilities are largely orthogonal^76–78^. Causal evidence that major interoceptive hub regions (e.g., the insula) encode cardiac afferents to regulate emotional states has been reported^79^, but it still remains an open question whether other modalities yield comparable mediating relationships or not, and which is most informative for inferring affective updating. Fourth, our spatial inference about the neural embeddings should be interpreted with caution. While our feature attribution converges on frontocentral delta oscillations previously linked to cardioception, the spatial resolution of our scalp EEG and its inherent physiological artifacts do not allow direct confirmation. Future studies employing high-density EEG with source localization or concurrent fMRI would enable a direct test of whether established interoceptive hub regions contribute to this representation. Last, our sample size for neurophysiological analyses may limit statistical power, particularly for detecting modest mediation effects and proving generalizability. Although we cross-validated using leave-one-subject-out procedures, independent replication in larger and clinically diagnosed cohorts will be important to establish the robustness and clinical generalizability.

In sum, we investigated whether, how, and when depressive symptom severity relates to delayed updating of affective states, and traced this delay to a speed-specific slowing of cardiac-coupled cortical dynamics. By combining a naturalistic movie-watching paradigm with the interoceptive account of affect and a dynamical-systems framework, we tested the embodied account of affective updating in a contextualized manner that context-agnostic approaches cannot achieve. Theoretically, this extends the Emotion Context Insensitivity framework from blunted response magnitude into a temporal dimension. Methodologically, our trajectory-based framework is modality- and disorder-agnostic, offering a general approach for quantifying brain-body dynamics across affective dysfunctions and beyond depression.

## Methods

### Participants

A total of forty-five healthy young adults (Korean; 27 females; *M*_age_ = 22.5 years, *SD*_age_ = 2.5 years) completed the experimental procedure. We excluded data from two participants: one identified as an outlier for baseline positive mood (exceeding 1.5 × interquartile range) and another due to a technical failure in recording continuous affect ratings. Consequently, data from forty-three participants were included in the behavioral analyses (*N* = 43). All participants met the following inclusion criteria: (i) no current use of psychiatric medications, (ii) no history of neurological or psychiatric disorders, and (iii) right-handedness. The study protocol was approved by the Institutional Review Board of Seoul National University, and all participants provided written informed consent prior to the experiment.

### Experimental paradigm

The experimental session lasted approximately 60mins and was conducted in a sound-attenuated room at the XR Experience Center, Seoul National University. The procedure consisted of five sequential phases: psychological assessment, resting-state recording, practice session, main movie-watching session, and post-movie questionnaire.

First, during the psychological assessment, participants provided demographic information and completed several validated Korean versions of psychometric scales via an online survey. These included the Positive and Negative Affect Schedule^80^ (K-PANAS) for baseline mood, the Beck Depression Inventory-II^81^ (K-BDI-II), and the Generalized Anxiety Disorder-7^82^ (K-GAD-7).

Second, resting-state signals were recorded for 2min using the Enobio 20 system (Neuroelectrics). This session aimed to evaluate signal quality and establish baseline neurophysiological signals for subsequent normalization. EEG data were recorded from 19 channels (F7, AF3, AF4, F8, F3, Fz, F4, FC5, FC6, C3, Cz, C4, CP5, CP6, P3, Pz, P4, PO3, and PO4) and a single-channel ECG (attached to the left rib) at a sampling rate of 500 Hz. Signal quality was monitored in real-time using the manufacturer’s embedded software, and a minimum of 30 sec of stable ‘good’ or ‘medium’ quality signals was ensured before proceeding.

Third, a practice session was conducted to familiarize participants with the real-time affect rating task using the Evaluative Space Grid^38^. The Evaluative Space Grid is a 2D plane where positive and negative affect are rated on independent 0–5 scales. A midpoint of 2.5 was used to categorize affective states into neutral, positive, negative, and mixed states (see ‘Behavioral analyses’), while allowing for parametric encoding of intensity within each quadrant. Participants were instructed to report their own subjective feelings rather than the emotions depicted in the movie. For the practice, we used a 4-min trailer of the movie *Up*, well-known for evoking rapid shifts in diverse affects. Real-time ratings were recorded by tracking the mouse cursor coordinates at 10Hz. Following the practice, all participants confirmed via ad-hoc questionnaires that the rating task did not interfere with their movie-watching experience and that the grid cells were clearly understood. EEG and ECG were not recorded during this session.

Fourth, during the main movie-watching session, participants viewed the 7-min animation *One Small Step*^39^. Concurrent with viewing, participants provided continuous affect ratings as practiced, and EEG/ECG signals were recorded following the same protocol as the resting-state session.

Finally, in the post-movie questionnaire, participants reported their global valence and arousal for the entire movie. Valence was assessed using a single cell on the Evaluative Space Grid, and arousal was measured on a 9-point Likert scale^83^. All participants reported no prior familiarity with *One Small Step*.

### Data Preprocessing and feature extraction

#### Affect ratings preprocessing

Real-time affect ratings collected during movie watching were preprocessed in three sequential steps. First, positive and negative affect ratings were downsampled from 10Hz to 2Hz (i.e., 0.5s intervals) to synchronize with the EEG feature time-series derived from short-time Fourier transform. Second, a Savitzky-Golay filter (window length = 25s; polynomial order = 2) was applied to both ratings to attenuate high-frequency noise while preserving the underlying affect dynamics. The Savitzky-Golay filter is particularly effective in filtering out high-frequency artifacts caused by sudden movements^84^ while accurately detecting low-frequency shifts in affect dynamics^85^. Filtering was implemented using the ‘scipy’ library^86^ (version 1.17.1). Third, any values exceeding the [0, 5] scale of the Evaluative Space Grid as a result of filtering were clipped to ensure they remained within the valid rating range.

#### EEG preprocessing

EEG signals were preprocessed using an automated pipeline based on EEGLAB (version 2025.0.0), validated to best preserve ground-truth neural signals among several algorithms^87^. First, the initial and final 3s of each recording were removed to rule out potential onset and offset effects (e.g., surprising effects or loss of attention). Second, a 0.5Hz high-pass filter was applied to the data. Third, poor-quality channels were identified using Artifact Subspace Reconstruction and replaced via spherical spline interpolation. Notably, as the CP6 channel exhibited poor signal quality in a majority of participants (*N* = 27/43), this channel was excluded from all participants, and the remaining 18 EEG channels were used for subsequent analyses. Finally, Independent Component Analysis was performed for artifact rejection. Components with a > 90% probability of being noise components (e.g., eye movements, head motion, or cardiac artifacts) were automatically identified and removed using the ICLabel plugin. Following automated preprocessing, all cleaned signals underwent rigorous visual inspection. Thirteen participants were excluded due to persistent poor signal quality, resulting in a final sample of thirty participants (*N* = 30) for all subsequent EEG and ECG analyses.

#### EEG feature extraction

To extract time-varying frequency-domain features, we applied short-time Fourier transform (STFT) to the preprocessed EEG signals across five frequency bands: delta (1-4Hz), theta (4-8Hz), alpha (8-14Hz), beta (14-31Hz), and gamma (31-49Hz). We used a 500-sample Hanning window (1,000ms) with a 250-sample hop size (500ms), resulting in 90 EEG features (18 channels × 5 frequency bands) at 0.5s intervals. Finally, to account for individual differences in baseline power, each feature was baseline-normalized by subtracting the feature-specific mean power calculated from the corresponding participant’s resting-state recording. STFT was implemented using the ‘scipy’ library^86^ (version 1.17.1).

#### ECG preprocessing

ECG signals from the thirty participants were preprocessed using the ‘NeuroKit2’ library^88^ (version 0.2.13). We leveraged NeuroKit2 due to its specialized pipeline for single-channel ECG preprocessing and its provision of standardized signal quality metrics. Specifically, we implemented the ‘elgendi2010’ pipeline^89^ for signal cleaning and R-peak detection. Following preprocessing, signal quality was evaluated based on established criteria^90^, and all participants exhibited “excellent” - the best signal quality across the entire recording.

#### ECG feature extraction

Following preprocessing, continuous beats-per-minute (BPM) were calculated based on the R-R intervals derived from the identified R-peaks. To ensure temporal synchronization with the STFT-derived EEG features, we applied a sliding window (window size = 500 samples, 1,000ms; hop size = 250 samples, 500ms) to compute the mean BPM within each window, yielding a smoothed BPM time-series at 0.5s intervals. Consequently, this pipeline produced a temporally aligned time-series of self-reported affect ratings (positive and negative feelings), EEG spectral features, and BPM for each participant at a common 0.5s resolution.

### Behavioral analysis

#### Identification of contextual transitions in movie

To identify the movie’s contextual transition timepoints based on group-level affective responses while avoiding circularity, we recruited 18 independent healthy young adults as annotators (Korean; 10 females; *M*_age_ = 20.1 years, *SD*_age_ = 1.5 years). Annotators followed the same protocol as the main experimental participants: they practiced real-time affect rating using the Evaluative Space Grid during the trailer of *Up* and provided continuous ratings while watching *One Small Step*. Their rating time-series were preprocessed identically to the main dataset (see ‘Affect ratings preprocessing’). We then calculated a group-level ‘normative trajectory’ by averaging the positive and negative affect ratings respectively across annotators at each timepoint. Afterward, we applied the ‘ruptures’ library^91^ (version 1.0.6) to this normative trajectory to detect abrupt shifts in affective states. Specifically, we employed a binary segmentation model, which has demonstrated superior performance in identifying change points within bivariate affective time-series^85^. Given the movie’s clear narrative structure of introduction-development-turn-conclusion, we constrained the model to identify three transition timepoints, which demarcated the movie into four major affective phases.

#### Calculation of affective states transition latency

For each of the three transition timepoints identified from the independent annotators’ ratings, we calculated the latency for participants in the main experiment to first enter context-congruent affective states. Based on the annotator data, the ‘congruent’ affective states for four movie phases were defined as neutral, positive, negative, and mixed, respectively (**Supplementary Fig 1**). We confirmed that these dominant states were also observed in the real-time affect ratings of the main participants (**Fig 1c and d**). To categorize each participant’s affective state at every timepoint, we applied a threshold of 2.5 (the midpoint of each rating axis) to the positive (POS) and negative (NEG) ratings as follows: neutral (POS < 2.5 and NEG < 2.5), positive (POS ≥ 2.5 and NEG < 2.5), negative (POS < 2.5 and NEG ≥ 2.5), and mixed (POS ≥ 2.5 and NEG ≥ 2.5). From the resulting affective state sequence for every participant, transition latency (in seconds) was defined as the time elapsed from a phase transition until the participant first entered the corresponding phase-dominant state (**Fig 2a**). Specifically, for transitions into Phases 2 and 3, we measured the time to reach positive and negative states, respectively. For the transition into Phase 4, we calculated the latency to enter either positive or mixed states. This approach accounted for the nuanced affective context of Phase 4 (i.e., achievement after grief), where the resultant state could manifest as either a purely positive state (reflecting the resolution of grief) or a mixed state (reflecting the co-occurrence of joy and lingering sadness). Consequently, three transition latencies were obtained for each participant. In a single case where a participant failed to enter a negative state upon transitioning to phase 3, the latency was calculated as the full duration of the phase 3.

#### Intraclass correlation estimation

To assess the consistency of subject-level latency rankings across the three movie phase transitions, we computed a two-way mixed-effects, single-measure consistency intraclass correlation coefficient (i.e., ICC(3, 1)) on per-subject phase latencies (in seconds), with subjects treated as targets and phases as raters. The coefficient was estimated from a two-way ANOVA partitioning between-subject and residual variance. Statistical significance was evaluated using the corresponding *F*-test, and 95% confidence intervals were derived analytically from *F*-distribution quantiles. This procedure was repeated across all discretization thresholds for discretization threshold parameters ranging from 2.25 to 2.75 in 0.05 increments.

#### Linear mixed effects modeling

We fitted a linear mixed-effects model to evaluate whether participants’ depressive symptoms, as measured by BDI-II total scores, significantly explained the variance in affective transition latency. Biological sex, age, transition type, and peak intensities of context-congruent affect were included as fixed-effect covariates. Since affect dynamics and magnitude are closely entangled, the affect intensity should be controlled to test the specific relationship between affective transition and psychiatric conditions^9,92–94^. For every participant, we calculated peak rating values of phase-congruent states for the phase 2, 3, and 4, and controlled them in our linear mixed model. Since a model with participant-level random slopes were not converged, we used the model with only random intercepts to account for individual baseline differences. Additionally, considering the skewed distribution of latency, we used the log-transformed latency, *y*\*, as the outcome variable as follows:

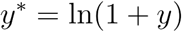

where *y* indicates a latency measure in seconds. Categorical inputs (i.e., sex and transition type) were dummy-coded, while continuous inputs (i.e., age, peak intensities, and BDI-II total score) were *z*-transformed prior to model fitting. In particular, peak intensities were normalized within participants. We confirmed that all key model assumptions (i.e., residual normality, homoscedasticity, linearity, and random intercept normality) were satisfied (**Supplementary Fig 3**). All statistical modeling was performed using the ‘statsmodels’ library^95^ (version 0.14.6).

#### Bayes factor approximation

To more directly evaluate the explanatory contribution of the BDI-II total score and the GAD-7 total score to the latency of affective transition, we computed Bayes factors for each regressor in addition to statistical significance. To this end, we applied a BIC-based Bayes factor approximation technique^96^. Specifically, the Bayes factor for the fixed effect of each regressor was defined as the natural exponential of the difference in Bayesian Information Criterion (BIC) between the ‘main’ model including the regressor and a ‘reference’ model including only covariates and a random intercept without the regressor, as follows:

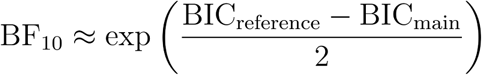

#### Model comparison for BDI-transition interaction

To test whether the association between BDI-II scores and affective transition latency was modulated by the valence of the changed contexts, we calculated the Bayes factor for the interaction between BDI-II scores and transition type. We fitted an expanded linear mixed-effects model adding the interaction term and compared its BIC with that of the main effects model to approximate the Bayes factor.

#### Leave-one-subject-out cross-validation

Predictive generalization of the BDI-II main-effects model was assessed at each discretization threshold via leave-one-subject-out (LOSO) cross-validation. For each held-out subject, the linear mixed effect model was refitted on the remaining 42 participants, and the held-out subject’s three phase log-latencies were predicted using fixed effects only (i.e., random intercept set to zero). Across all 43 folds, the 129 predicted-observed pairs were pooled, and *R*-squared was reported for every threshold parameter. To prevent a data leakage, *z*-transformation was performed within the training folds and held-out sample’s input features were scaled using training samples’ statistics.

### Modeling and validation of cardiac-coupled cortical representations

#### CEBRA-Behavior model fitting

To identify a latent cortical representation coupled to cardiac states, we fitted CEBRA-Behavior models to the EEG and *z*-transformed BPM time-series. BPM was selected as the auxiliary variable because it provides a sample-by-sample index of cardiac state that matches the second-by-second resolution of our analytic framework, an approach widely adopted in naturalistic neuroimaging studies (e.g., the ref.^32,97^). In contrast, other measures, for example, heart rate variability indices generally require longer analysis windows and reflect autonomic regulation operating over different timescales^98^. CEBRA-Behavior utilizes contrastive learning with minimizing infoNCE loss to align temporally synchronized EEG and BPM time-series, positioning EEG samples with similar BPM values closer in the latent space while repelling those with disparate values. Multi-session training was employed to project embeddings from all participants into a shared latent space. To determine the optimal dimensionality, we systematically explored dimensions ranging from 2 to 9. The 9-dimensional model was selected for subsequent analyses as it achieved the highest contrastive performance, defined as the average InfoNCE loss over the final 50 of 1,000 iterations. A 9-dimensional null model was also fitted using the EEG time-series and randomly shuffled BPM data. By doing so, we extracted each participant’s best and null latent embeddings from the best and null model, respectively. Hyperparameters were held constant across all models: model_architecture = ‘offset10-model’, max_iterations = 1,000, learning_rate = 0.001, num_hidden_units = 90 (number of EEG features), batch_size = 863 (number of timepoints), and time_offsets = 2 (1 sec). All modeling was conducted using the ‘cebra’ library^41^ (version 0.5.0).

#### Consistency evaluation

We evaluated whether each participant’s cardiac-conditioned cortical representations exhibited a reliable geometric structure both within and across individuals. Reliability was quantified using within- and between-participants consistency scores, which were compared against scores derived from null models. In CEBRA, the consistency score is defined as the *R*-squared value of a linear regression model between two embeddings, representing the degree to which they can be aligned via linear mapping^41^. First, to assess within-participants consistency, we calculated representative consistency scores for each participant. For this purpose, we trained four additional 9-dimensional multi-session models using identical datasets and hyperparameters, resulting in five models in total. For each participant, a representative consistency score was obtained by averaging the 20 possible pairwise consistency scores among the five embeddings. Second, to evaluate between-participants consistency, we computed representative consistency scores using the original 9-dimensional model. For each participant, the representative score was calculated as the average consistency score across all pairs involving that participant and the other 29 participants. For both metrics, statistical significance was tested using subject-wise paired two-sided *t*-tests against the consistency scores derived from the null models.

#### Biological plausibility evaluation

To evaluate the biological plausibility of the modeled cortical representation, we investigated which EEG features were most instrumental in constructing the cardiac-coupled space using a feature-ablation-based decoding analysis. We quantified feature importance by measuring the decrease in decoding performance. Specifically, we measured how much the accuracy of predicting BPM from the resultant embeddings dropped when individual EEG features were ablated.

For each participant, we first established a reference performance score through a four-step decoding procedure: (i) extracting the 9-dimensional embeddings from the full EEG time-series using the best CEBRA model, (ii) splitting the time-series into training and test sets using a 7:3 chronological split, (iii) training a *k*-nearest neighbors (kNN) regressor to predict BPM from the training embeddings, and (iv) calculating the *R*-squared between the predicted and actual BPM on the test set.

Subsequently, we iteratively ablated each of the 90 EEG features by replacing its values with the participant-specific time-series mean and recalculated the decoding performance. Feature importance for each feature was then defined as the effect size (Cohen’s *d*) representing the difference between the reference and ablated performance scores across all participants. All decoding analyses were performed using the ‘scikit-learn’ library^99^ (version 1.8.0).

### Cardiac-coupled cortical dynamics analysis

#### Calculation of dynamics features

From the 9-dimensional latent embeddings extracted using the best CEBRA-Behavior model, we quantified the embedding dynamics at each contextual transition timepoint in terms of speed and angle^30,31^. To this end, we defined a ‘transition window’, spanning [−*k*, +*k*] around each contextual transition, with a window size parameter *k* in seconds. Given that CEBRA-Behavior embeddings are projected onto a unit hypersphere, *S*^8^, rather than Euclidean space, we employed Riemannian geometry to calculate speed and angle for all timepoints within the transition windows.

First, the speed *θ_t_* of a participant’s brain-heart embedding *Z* at time *t* was defined as the geodesic distance between *Z_t_* and *Z_t_*_+1_, calculated as follows:

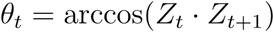

Second, the angle *ϕ_t_* at time *t* was defined as the directional change between consecutive tangent vectors. To compute this, we first obtained the tangent vector, *v_t_* pointing from *Z_t_*and *Z_t_*_+1_ in the tangent space via the logarithmic map:

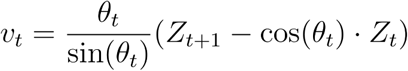

Because *v_t_* and *v_t+_*_1_ reside in distinct tangent spaces, they cannot be compared directly. Therefore, *v_t_* was parallel-transported to the tangent space of *v_t+_*_1_ to obtain *v*\**_t_*:

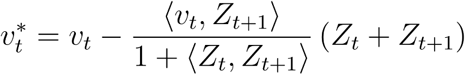

Based on this, the final angle *ϕ_t_* was defined as the angle between the parallel-transported vector *v*\**_t_* and the subsequent tangent vector *v_t+_*_1_:

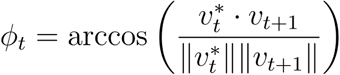

In this way, *θ* and *ϕ* values for all timepoints within the three transition windows were calculated for each participant. To evaluate the within-participant reliability of the speed and angle features across the three phase transitions, we computed the intraclass correlation for each feature (see ‘intraclass correlation estimation’) and confirmed significant and modest reliability for both dynamic features. Finally, for each participant, the speed and angle features were respectively averaged across transitions for each of the three transitions to derive representative speed and angle features. For all primary analyses, the default window size was set to *k* = 5; however, to demonstrate that our findings were robust to this parameter, we repeated all analyses while increasing the window size from 3 to 7s in 0.5s increments.

#### Mediation model testing

We performed mediation analyses to test whether brain-heart dynamics (speed or angle) mediate the relationship between depressive states and delayed latency of affective state transition. Specifically, we fitted separate mediation models for each mediator of embedding speed and angle with BDI-II total scores as the input, brain-heart dynamic feature as the mediator, and the averaged log-transformed latency of affective transitions as the outcome. In all models, age, dummy-coded sex, and phase-averaged peak intensities were included as covariates to control for their potential confoundings. Except for biological sex, all continuous features were *z*-scored prior to analysis. For all paths except the indirect path, beta coefficients, 95% confidence intervals (CI), and *P*-values were estimated using ordinary least squares regression. For the indirect path, 95% CIs and *P*-values were calculated using 10,000 bootstrap iterations in a two-sided manner. To ensure that the mediating effects of brain-heart speed and angle were robust to the choice of transition window size, we tested a total of 18 models (2 mediators × 9 window sizes), with window size parameters (*k*) ranging from 3 to 7s in 0.5s increments. All mediation analyses were implemented using the ‘pingouin’ library^100^ (version 0.5.5).

#### Leave-one-subject-out cross-validation

For the mediation model, we performed LOSO cross-validation tests at each window size and for each mediator. Because data were subject-level unlike in the linear mixed effects model of behavioral analysis (see ‘Behavioral analyses’), the held-out subject’s mean log-latency was predicted from an OLS regression on BDI-II total score, mediator, and covariates fitted to the remaining 29 participants. *R*-squared were computed across the 30 prediction-observation pairs per window size and mediator combination. To prevent data leakage, all normalizations were performed within training folds.

#### Model comparison using permutation

We evaluated the statistical differences of predictive performances among the speed-mediator, angle-mediator, and base models that include only covariates. Models were compared by their held-out *R*-squared with the pairwise difference, *ΔR*^2^, as the test statistic. For each pairwise, null distributions were built from 10,000 permutations. On each permutation, only the mediator column of a given model (i.e., speed or angle) was reshuffled across subjects, leaving the outcome, the BDI-II score and the covariate block (i.e., age, sex, phase-averaged peak affect intensity) intact, and the entire LOSO-CV procedure was re-run on the permuted data. Permuting a mediator rather than the outcome matches model dimension exactly between observed and permuted fits, so both the variance explained by the remaining predictors and the out-of-sample cost of the additional parameters are still represented in the null. Under the hypothesis that the *R*-squared would follow the order of speed-mediator > angle-mediator > base model, one-sided *P*-values, *p*_perm_, were computed as follows:

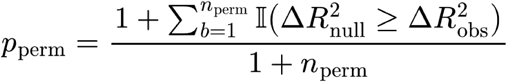

#### Control analyses

We evaluated the specificity of the mediation model involving BDI-II scores, brain-heart dynamic features, and affective transition latency through three distinct control analyses. First, we tested a mediation model using GAD-7 total scores that represent anxiety severity as the input variable, investigating whether the observed effects were specific to depressive states. Second, to assess modality specificity, we trained a CEBRA-Time model, which performs contrastive learning based on timepoint indices rather than cardiac signals, to test whether purely neural latent embeddings mediate the relationship. For CEBRA-Time, we compared contrastive performance across 2-9 dimensions using the same hyperparameters as the CEBRA-Behavior model and selected the 5-dimensional model as the best fit (**Supplementary Fig 6**). Using this control model, we calculated embedding speed and angle in the same manner and tested their indirect effects in the mediation model. In addition to the 5-dimensional model (**Fig 5b**), which was identified as the optimal dimensionality for the CEBRA-Time model trained using EEG alone without ECG, we performed the same control analysis using a 9-dimensional embedding matching the dimensionality of the CEBRA-Behavior model used in the main analysis (**Supplementary Fig 7**). Last, to examine temporal specificity, we calculated embedding speed and angle from ‘control windows’ centered at the midpoints of the first three movie phases and used them as mediators to verify whether the mediation was specific to the moments of contextual transition. As in the main analysis, all control models were evaluated across 9 window size parameters ranging from 3 to 7s in 0.5-s increments, resulting in a total of 54 control mediation models (2 mediators × 3 control types × 9 window sizes).

## Acknowledgement

This work was supported by the National Research Foundation of Korea (NRF) grant funded by the Korea government (MSIT) (No. 2021R1C1C1006503, RS-2023-00266787, RS-2023-00265406, RS-2024-00421268, RS-2024-00342301, RS-2024-00435727, RS-2025-25457239, RS-2026-25490183, RS-2026-25524667, RS-2026-25518389, RS-2021-NR061370, NRF-2021M3E5D2A01022515, and NRF-2021S1A3A2A02090597), by the Researchers Program through Seoul National University (No. 200-20250071, 200-20250049, 200-20240057, 0670-20260027, 200-20260009, 200-20250116, 200-20260081, 200-20250115, 200-20250113, 0670-20250039, 200-20260009). Additional support was provided by the Institute of Information & Communications Technology Planning & Evaluation (IITP) grant funded by the Korea government (MSIT) [No. RS-2021-II211343, Artificial Intelligence Graduate School Program, Seoul National University] and by the Global Research Support Program in the Digital Field (RS-2024-00421268). This work was also supported by the Artificial Intelligence Industrial Convergence Cluster Development Project funded by the Ministry of Science and ICT and Gwangju Metropolitan City, by the Korea Brain Research Institute (KBRI) basic research program (25-BR-05-01), by the Korea Health Industry Development Institute (KHIDI) and the Ministry of Health and Welfare, Republic of Korea (HR22C1605), and by the Korea Basic Science Institute (National Research Facilities and Equipment Center) grant funded by the Ministry of Education (RS-2024-00435727). We acknowledge the National Supercomputing Center for providing supercomputing resources and technical support (KSC-2023-CRE-0568, KSC-2024-CRE-0198, KSC-2025-CRE-0340).

## Code/Data Availability

All code and data required to reproduce and visualize all results of this study will be made publicly available through GitHub and Zenodo following publication.

## Author Contributions

**J.L.:** conceptualization, data collection, formal analysis, investigation, methodology, project administration, visualization, writing-original draft, and writing-review & editing. **K.O.:** data collection, investigation, and writing-review & editing. **J.K.:** investigation and writing-review & editing. **J.C.:** conceptualization, funding acquisition, investigation, supervision, and writing-review & editing.

---

**Supplementary Fig 1.**
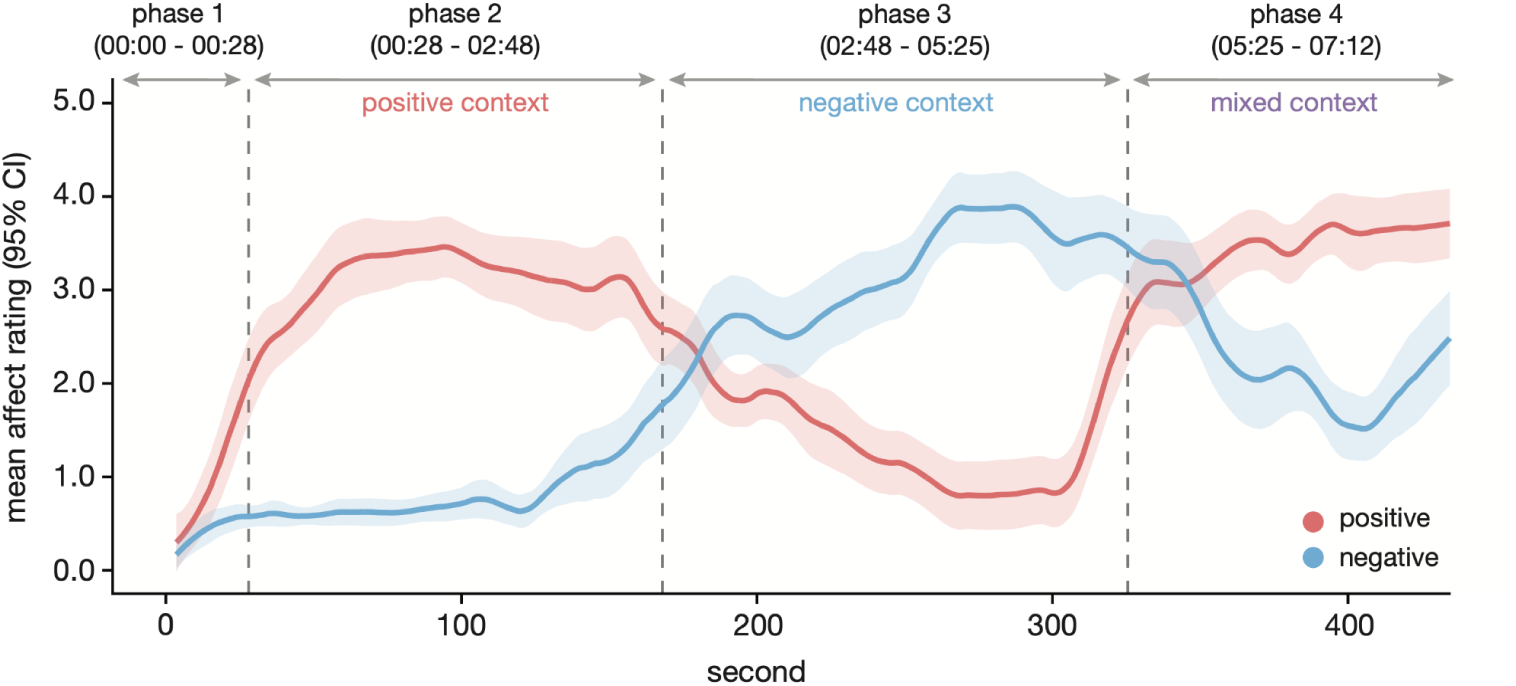
Group-level dynamics of independent annotators’ affect ratings. Mean and 95% confidence intervals (CIs) of continuous affect ratings across timepoints, as reported by independent annotators (*N* = 18) while viewing *One Small Step*. Grey vertical dashed lines indicate three affective context transitions identified by applying change-point detection algorithms to the group-level affect dynamics (see ‘Identification of contextual transitions in movie’ section in **Methods**).

**Supplementary Fig 2.**
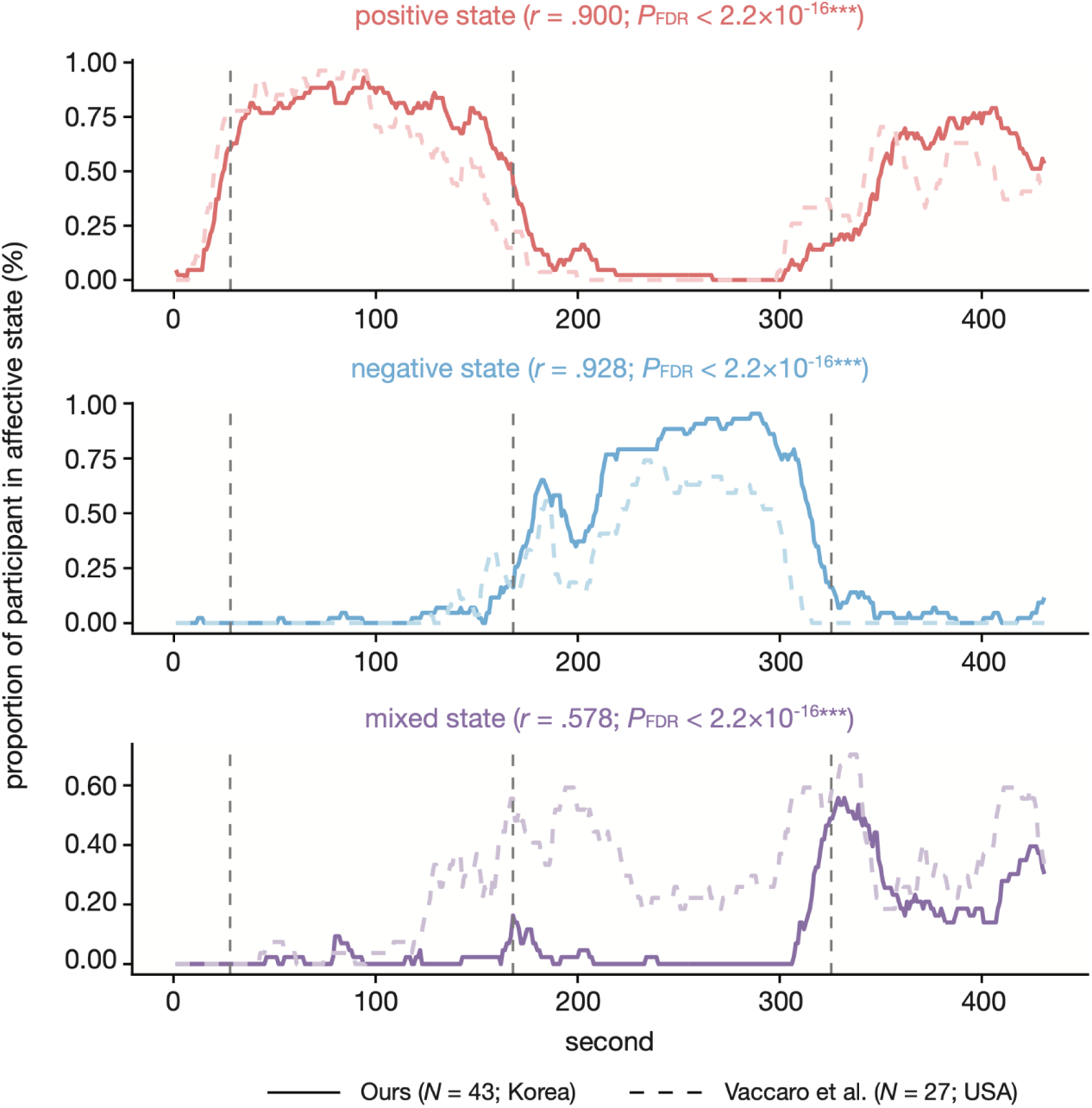
Temporal similarities of positive, negative, and mixed affect dynamics between Vaccaro et al. (2024) and ours. Temporal patterns of affect ratings were compared between the current study and the ref.1. In ref.^1^, participants (*N* = 27) reported their affective states categorically (positive, negative, mixed, or neutral) via continuous button presses while watching the same movie *One Small Step*. To align the datasets, continuous ratings from the current study’s Evaluative Space Grid (*N* = 43) were converted into affective state sequences using a threshold of 2.5. Group-level dynamics were then derived by calculating the proportion of participants in each state at every timepoint for both studies. For statistical comparison, the current study’s data were downsampled from 0.5s to 1.0s intervals to match the temporal resolution of Vaccaro et al. Pearson’s correlation coefficients were calculated for each affective state to assess temporal similarity. Grey vertical dashed lines indicate phase transition timepoints. We confirm that participants in both studies display highly aligned temporal patterns in their group-level affect dynamics despite the differences in cultural background and self-report paradigm. All *P*-values were False Discovery Rate (FDR) corrected; \**P*_FDR_ < .05; \*\**P*_FDR_ < .01; \*\*\**P*_FDR_ < .001.

**Supplementary Fig 3.**
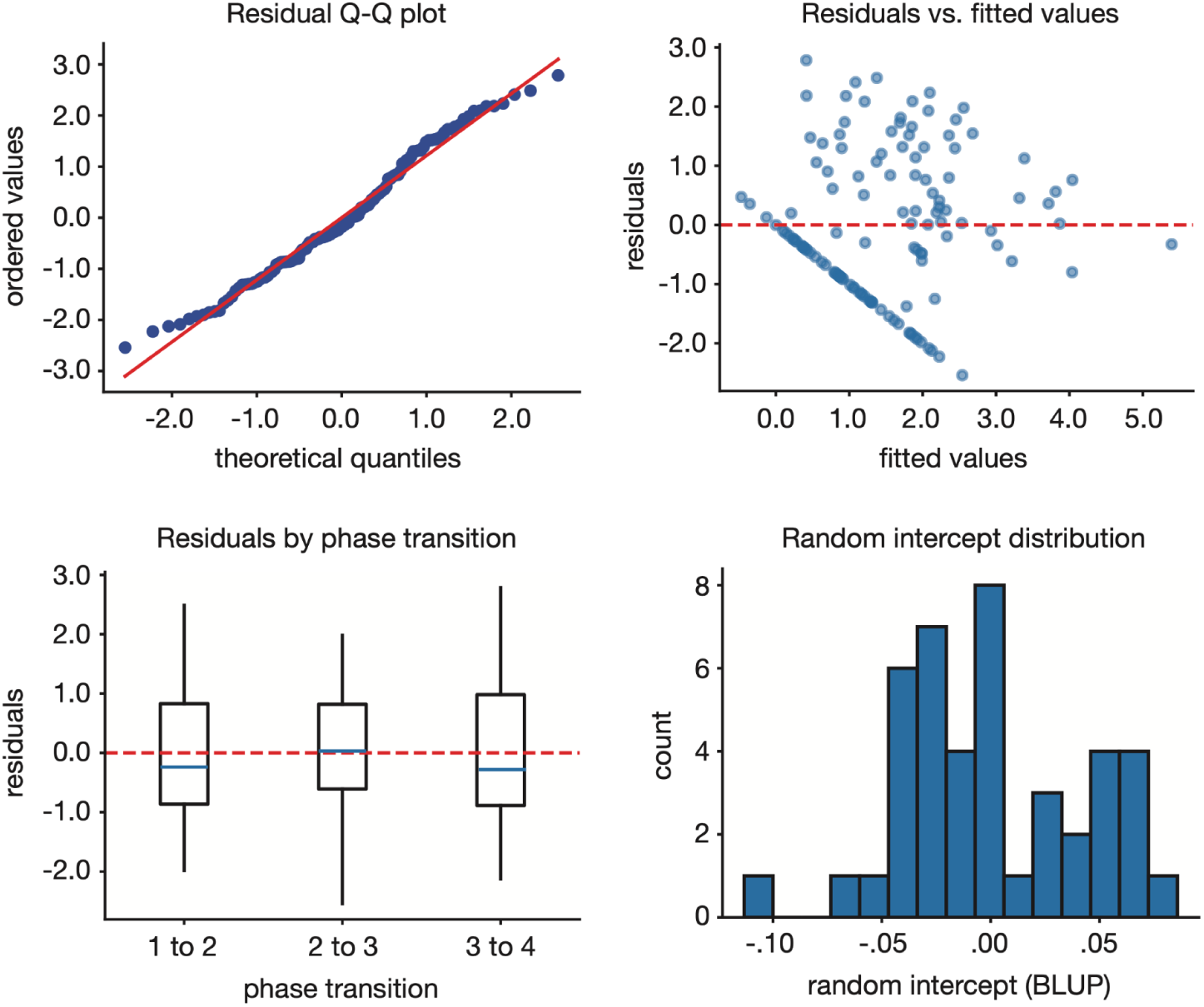
Assumption check for linear mixed effects modeling. We evaluated four fundamental assumptions of the linear mixed effects model, which included BDI-II as a fixed effect and subject-specific random intercepts: normality of residuals, linearity, homoscedasticity, and normality of random intercepts. **(Top left)** The Residual Q-Q plot visually assesses the normality of residuals; the alignment of residuals with the red diagonal reference line indicates that the normality assumption was largely satisfied. Shapiro-Wilk test revealed that the normality assumption could not be rejected (*W* = .980, *P* = .057). **(Top right)** To evaluate homoscedasticity and linearity, residuals were plotted against fitted values. The random distribution of residuals around zero, without discernible systematic patterns, confirms both assumptions. **(Bottom left)** Residual distributions across phase transition types show no significant variance differences, satisfying the assumption of homoscedasticity across experimental phases. The Levene test showed that equal residual variance across phases could not be rejected (*F* = .110, *P* = .896). **(Bottom right)** The distribution of random intercepts estimated via Best Linear Unbiased Predictors (BLUPs) was examined for normality. Shapiro-Wilk test confirmed that the normality assumption could not be rejected (*W* = .961, *P* = .151). Collectively, these diagnostics confirm that all model assumptions were not violated.

**Supplementary Fig 4.**
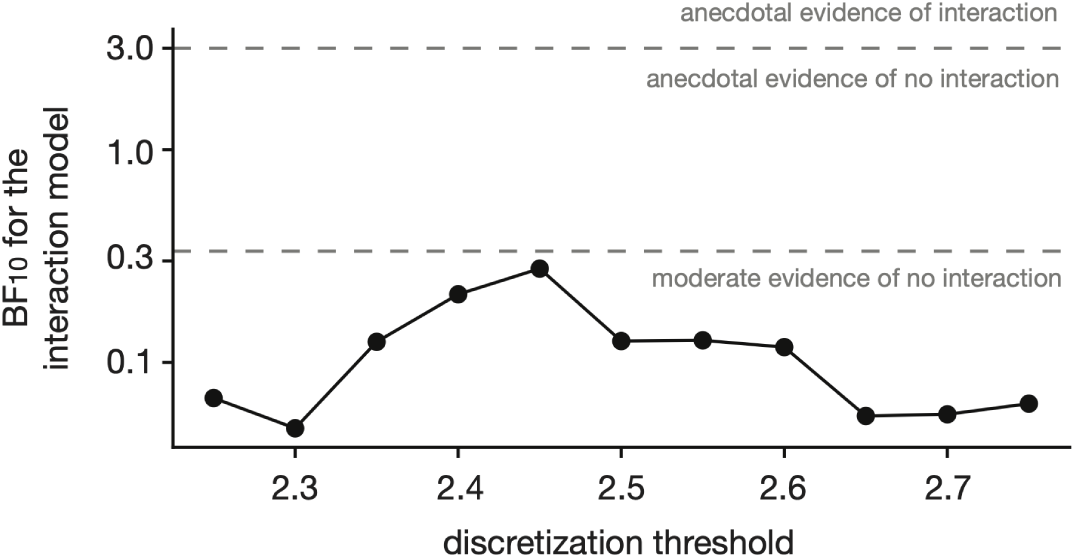
Bayes factors of the interaction between BDI-II total score and phase transition type across discretization thresholds. To examine whether the association between depressive states and affective transition latency is modulated by the valence of the transition phase, we calculated the Bayes factor (BF_10_) for the model including the interaction term between BDI-II total scores and phase transition types using a linear mixed-effects framework. Specifically, BF_10_ was approximated by comparing the Bayesian Information Criteria of a base model with a model containing the interaction term (see ‘Bayes factor approximation’ in **Methods**). We observed moderate evidence favoring the absence of an interaction, a result that remained robust across various affective state discretization thresholds. The interpretation of Bayes factors followed the criteria proposed by the ref.^2^.

**Supplementary Fig 5.**
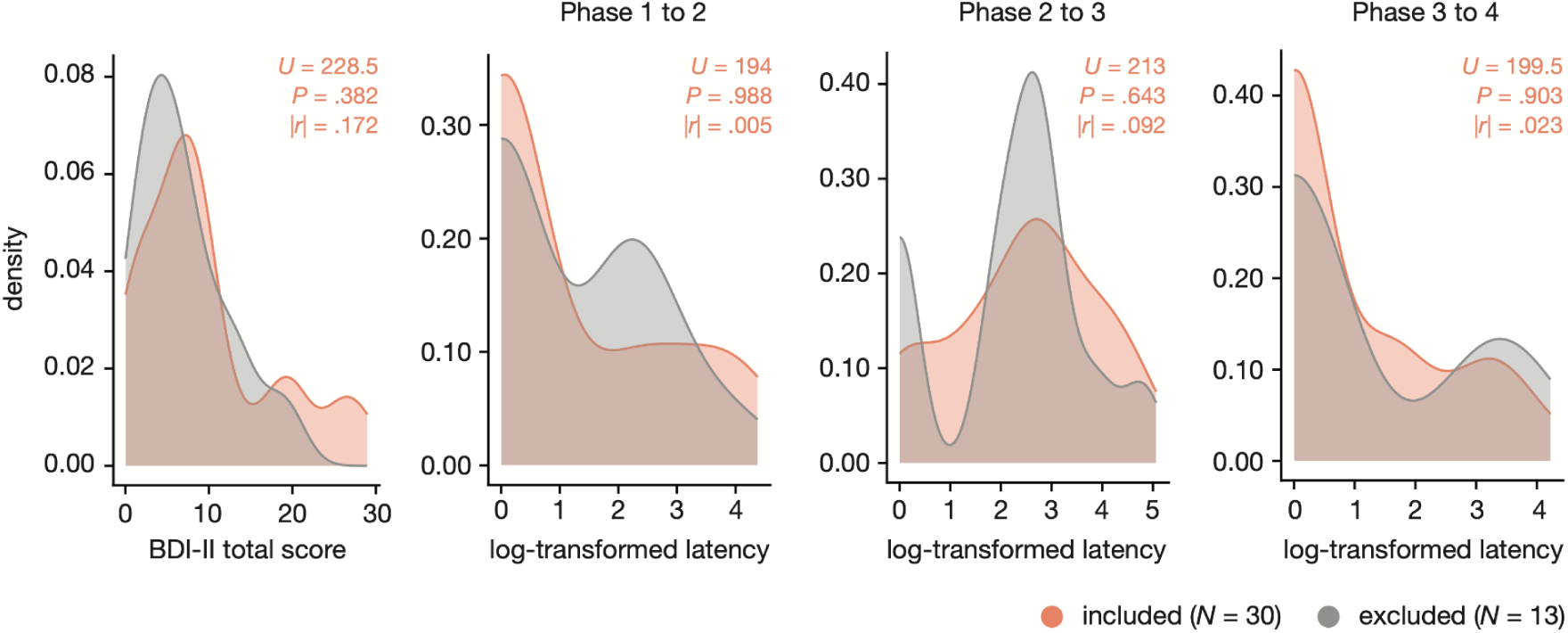
Confounding test of participant subsampling in neurophysiological analyses. To ensure that the exclusion of 13 participants due to neurophysiological signal quality did not introduce confounding bias into the mediation analysis, we compared BDI-II total scores and log-transformed transition latencies between the included and excluded groups. Differences were assessed using two-sided Mann-Whitney U tests, with effect sizes quantified as the absolute value of the rank-biserial correlation coefficient, |*r*|. No statistically significant differences were observed across any of the four variables at *α* = .05, and effect sizes were negligible. These results support the conclusion that the exclusion of these participants did not exert a substantial confounding effect on the primary findings.

**Supplementary Fig 6.**
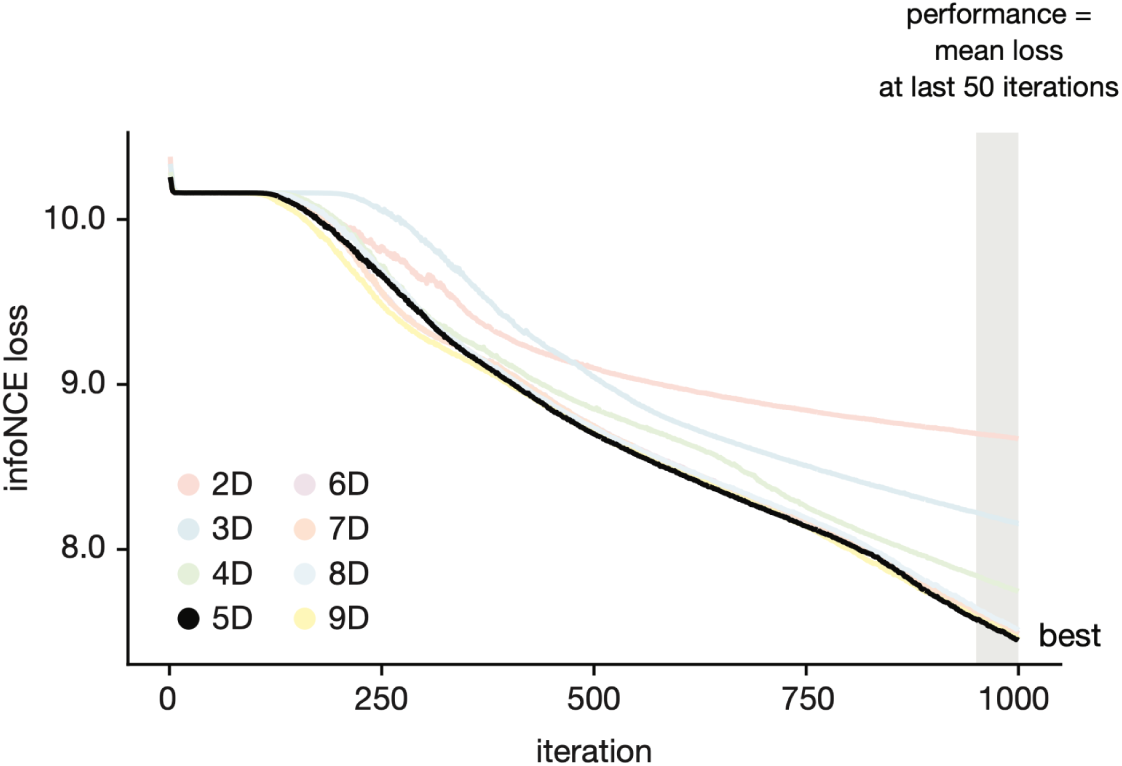
Dimensionality selection for CEBRA-Time model. Training curves for CEBRA-Time models across latent dimensions ranging from 2 to 9 are shown, using the timepoint index as an auxiliary variable. Consistent with the CEBRA-Behavior model that used beats per minute time-series as an auxiliary variable, the final performance was defined as the mean infoNCE loss averaged over the last 50 iterations. Based on this criterion, the 5-dimensional model was selected as the optimal model and used in the control analysis (see ‘Control analyses’ section in **Methods**).

**Supplementary Fig 7.**
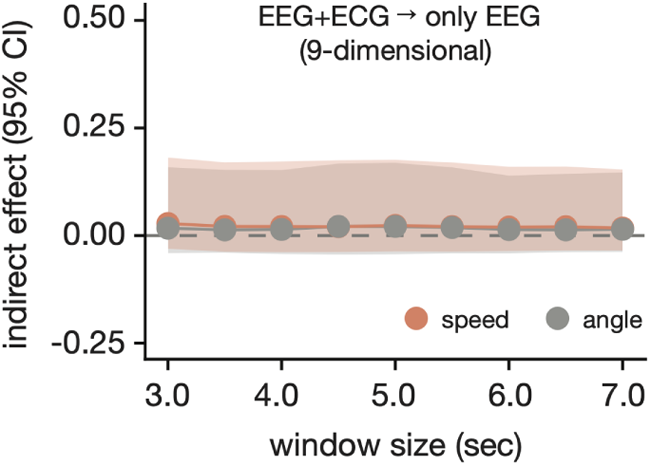
Indirect effects of neural dynamics without cardiac information under 9-dimensional latent space. In addition to the 5-dimensional model (see **Fig 5b**), which was identified as the optimal dimensionality for the CEBRA-Time model trained using EEG alone without ECG, we performed the same control analysis using a 9-dimensional embedding matching the dimensionality of the CEBRA-Behavior model used in the main analysis. All confidence intervals of indirect effects were estimated using 10,000 times bootstrapping for window sizes ranging from 3 to 7s in 0.5s increments.

**Supplementary Table 1.** Phase-wise comparisons of affective ratings and heart rate.

| Feature | Phase A | Phase B | *t*-value | *df* | $P_{\text{FDR}}$ | Hedges' *g* |
| --- | --- | --- | --- | --- | --- | --- |
| *Mean positive affect (N = 43)* |  |  |  |  |  |  |
|  | Phase 1 | Phase 2 | -15.513 | 42 | < 0.001*** | -2.971 |
|  | Phase 1 | Phase 3 | -0.401 | 42 | 0.690 | -0.087 |
|  | Phase 1 | Phase 4 | -13.858 | 42 | < 0.001*** | -2.997 |
|  | Phase 2 | Phase 3 | 19.959 | 42 | < 0.001*** | 4.030 |
|  | Phase 2 | Phase 4 | -1.636 | 42 | 0.131 | -0.198 |
|  | Phase 3 | Phase 4 | -19.282 | 42 | < 0.001*** | -3.908 |
| *Mean negative affect (N = 43)* |  |  |  |  |  |  |
|  | Phase 1 | Phase 2 | -9.875 | 42 | < 0.001*** | -1.167 |
|  | Phase 1 | Phase 3 | -30.166 | 42 | < 0.001*** | -5.256 |
|  | Phase 1 | Phase 4 | -14.600 | 42 | < 0.001*** | -2.646 |
|  | Phase 2 | Phase 3 | -23.339 | 42 | < 0.001*** | -4.254 |
|  | Phase 2 | Phase 4 | -9.868 | 42 | < 0.001*** | -1.841 |
|  | Phase 3 | Phase 4 | 7.882 | 42 | < 0.001*** | 1.582 |
| *Mean beats-per-minute (z-scored; N = 30)* |  |  |  |  |  |  |
|  | Phase 1 | Phase 2 | -0.391 | 29 | 0.863 | -0.090 |
|  | Phase 1 | Phase 3 | -0.175 | 29 | 0.863 | -0.048 |
|  | Phase 1 | Phase 4 | -3.999 | 29 | < 0.001*** | -1.185 |
|  | Phase 2 | Phase 3 | 0.285 | 29 | 0.863 | 0.090 |
|  | Phase 2 | Phase 4 | -5.939 | 29 | < 0.001*** | -1.902 |
|  | Phase 3 | Phase 4 | -7.429 | 29 | < 0.001*** | -2.095 |

*Note.* Statistical differences between two phases were tested with two-sided paired *t*-tests.
Beats-per-minute was z-scored within participants. All *P*-values were False Discovery Rate (FDR) corrected; \* $P_{\text{FDR}} < .05$ ; \*\* $P_{\text{FDR}} < .01$ ; \*\*\* $P_{\text{FDR}} < .001$ .

**Supplementary Table 2.** Reliability of affective transition latency across discretization thresholds.

| Threshold | ICC(3, 1) | F-value | $df_1$ | $df_2$ | P-value | Lower bound of 95% CI | Upper bound of 95% CI |
| --- | --- | --- | --- | --- | --- | --- | --- |
| 2.25 | 0.268 | 2.100 | 42 | 84 | 0.002** | 0.080 | 0.470 |
| 2.30 | 0.248 | 1.991 | 42 | 84 | 0.004** | 0.060 | 0.450 |
| 2.35 | 0.232 | 1.908 | 42 | 84 | 0.006** | 0.050 | 0.440 |
| 2.40 | 0.299 | 2.280 | 42 | 84 | 0.001** | 0.110 | 0.500 |
| 2.45 | 0.292 | 2.240 | 42 | 84 | 0.001** | 0.100 | 0.490 |
| 2.50 | 0.287 | 2.208 | 42 | 84 | 0.001** | 0.100 | 0.490 |
| 2.55 | 0.308 | 2.335 | 42 | 84 | < 0.001*** | 0.120 | 0.500 |
| 2.60 | 0.305 | 2.317 | 42 | 84 | 0.001** | 0.120 | 0.500 |
| 2.65 | 0.277 | 2.148 | 42 | 84 | 0.001** | 0.090 | 0.480 |
| 2.70 | 0.331 | 2.487 | 42 | 84 | < 0.001*** | 0.140 | 0.530 |
| 2.75 | 0.351 | 2.625 | 42 | 84 | < 0.001*** | 0.160 | 0.540 |

*Note.* Across the three movie phase transitions, we computed a two-way mixed-effects, single-measure consistency intraclass correlation coefficient - ICC(3, 1) on per-subject phase latencies (in seconds), with subjects treated as targets and phases as raters. The coefficient was estimated from a two-way ANOVA partitioning between-subject and residual variance. Statistical significance was evaluated using the corresponding *F*-test, and 95% confidence intervals were derived analytically from *F*-distribution quantiles; \**P* < .05; \*\**P* < .01; \*\*\**P* < .001.

**Supplementary Table 3.** Statistics of the linear mixed effect models across discretization thresholds.

| Threshold | Input variable | Beta coefficients | *P*-value | Lower bound of 95% CI | Upper bound of 95% CI |
| --- | --- | --- | --- | --- | --- |
| 2.25 | BDI-II | 0.219 | 0.060 | -0.009 | 0.446 |
|  | GAD-7 | -0.015 | 0.901 | -0.252 | 0.222 |
| 2.30 | **BDI-II** | **0.246** | **0.028\*** | **0.026** | **0.466** |
|  | GAD-7 | 0.006 | 0.958 | -0.226 | 0.239 |
| 2.35 | **BDI-II** | **0.279** | **0.005\*\*** | **0.084** | **0.474** |
|  | GAD-7 | 0.072 | 0.506 | -0.140 | 0.284 |
| 2.40 | **BDI-II** | **0.312** | **0.002\*\*** | **0.110** | **0.514** |
|  | GAD-7 | 0.065 | 0.560 | -0.153 | 0.282 |
| 2.45 | **BDI-II** | **0.314** | **0.003\*\*** | **0.109** | **0.520** |
|  | GAD-7 | 0.057 | 0.621 | -0.168 | 0.282 |
| 2.50 | **BDI-II** | **0.339** | **0.003\*\*** | **0.119** | **0.558** |
|  | GAD-7 | 0.034 | 0.783 | -0.209 | 0.277 |
| 2.55 | **BDI-II** | **0.313** | **0.009\*\*** | **0.080** | **0.546** |
|  | GAD-7 | -0.001 | 0.996 | -0.253 | 0.252 |
| 2.60 | **BDI-II** | **0.287** | **0.029\*** | **0.030** | **0.543** |
|  | GAD-7 | -0.014 | 0.917 | -0.286 | 0.257 |
| 2.65 | BDI-II | 0.262 | 0.051 | -0.001 | 0.525 |
|  | GAD-7 | -0.028 | 0.843 | -0.303 | 0.247 |
| 2.70 | BDI-II | 0.247 | 0.060 | -0.010 | 0.505 |
|  | GAD-7 | -0.037 | 0.788 | -0.305 | 0.231 |
| 2.75 | BDI-II | 0.256 | 0.059 | -0.009 | 0.522 |
|  | GAD-7 | -0.019 | 0.895 | -0.296 | 0.258 |

*Note.* The linear mixed effect models used the corresponding fixed regressor while controlling dummy coded sex, age, and phase transition type. The random intercept was included. All continuous variables were z-scored. Rows in bold indicate conditions where fixed effects are significant; \* $P < .05$ ; \*\* $P < .01$ ; \*\*\* $P < .001$ .

**Supplementary Table 4.** Reliability of brain-heart dynamics measures across window size parameters.

| Window size (sec) | Reliability of **speed** |  |  |  | Reliability of **angle** |  |  |  |
| --- | --- | --- | --- | --- | --- | --- | --- | --- |
|  | ICC(3,1) | P-value | Lower bound of 95% CI | Upper bound of 95% CI | ICC(3,1) | P-value | Lower bound of 95% CI | Upper bound of 95% CI |
| 3.0 | 0.259 | 0.010* | 0.040 | 0.500 | 0.212 | 0.028* | -0.010 | 0.460 |
| 3.5 | 0.291 | 0.005** | 0.070 | 0.530 | 0.321 | 0.002** | 0.100 | 0.560 |
| 4.0 | 0.320 | 0.002** | 0.100 | 0.550 | 0.347 | 0.001** | 0.120 | 0.580 |
| 4.5 | 0.331 | 0.002** | 0.110 | 0.560 | 0.396 | < .001*** | 0.170 | 0.620 |
| 5.0 | 0.323 | 0.002** | 0.100 | 0.560 | 0.402 | < .001*** | 0.180 | 0.620 |
| 5.5 | 0.343 | 0.001** | 0.120 | 0.570 | 0.346 | 0.001** | 0.120 | 0.580 |
| 6.0 | 0.332 | 0.002** | 0.110 | 0.560 | 0.366 | 0.001** | 0.140 | 0.590 |
| 6.5 | 0.353 | 0.001** | 0.130 | 0.580 | 0.311 | 0.003** | 0.090 | 0.550 |
| 7.0 | 0.377 | < .001*** | 0.150 | 0.600 | 0.260 | 0.010* | 0.040 | 0.500 |

*Note.* For each brain-heart dynamics feature, we computed a two-way mixed-effects, single-measure consistency intraclass correlation coefficient - ICC(3, 1) on per-subject phase latencies (in seconds), with subjects treated as targets and phases as raters. The coefficient was estimated from a two-way ANOVA partitioning between-subject and residual variance. Statistical significance was evaluated using the corresponding *F*-test, and 95% confidence intervals were derived analytically from *F*-distribution quantiles; \**P* < .05; \*\**P* < .01; \*\*\**P* < .001.

**Supplementary Table 5.** Indirect effect estimates of brain-heart dynamics mediators across time window parameters.

| Window size (sec) | Indirect effects of **speed** |  |  |  | Indirect effects of **angle** |  |  |  |
| --- | --- | --- | --- | --- | --- | --- | --- | --- |
|  | Beta coefficient | *P*-value | Lower bound of 95% CI | Upper bound of 95% CI | Beta Coefficient | *P*-value | Lower bound of 95% CI | Upper bound of 95% CI |
| 3.0 | **0.171** | **0.047\*** | **0.029** | **0.408** | 0.011 | 0.742 | -0.025 | 0.149 |
| 3.5 | **0.193** | **0.030\*** | **0.043** | **0.440** | 0.014 | 0.745 | -0.040 | 0.160 |
| 4.0 | **0.187** | **0.032\*** | **0.043** | **0.426** | 0.024 | 0.564 | -0.025 | 0.171 |
| 4.5 | **0.187** | **0.033\*** | **0.043** | **0.426** | 0.021 | 0.621 | -0.022 | 0.171 |
| 5.0 | **0.174** | **0.041\*** | **0.036** | **0.416** | 0.028 | 0.542 | -0.022 | 0.190 |
| 5.5 | **0.182** | **0.046\*** | **0.033** | **0.418** | 0.047 | 0.377 | -0.019 | 0.232 |
| 6.0 | **0.195** | **0.027\*** | **0.050** | **0.429** | 0.035 | 0.499 | -0.025 | 0.193 |
| 6.5 | **0.184** | **0.039\*** | **0.036** | **0.415** | 0.033 | 0.495 | -0.027 | 0.193 |
| 7.0 | **0.186** | **0.042\*** | **0.031** | **0.414** | 0.022 | 0.619 | -0.028 | 0.172 |

*Note.* The mediation model used the BDI-II total score as the input variable, the phase-averaged log-transformed latency measure as the output variable, and each dynamic feature as a mediator. Dummy-coded sex, age, and phase-averaged peak intensity of phase-congruent affect were controlled as covariates. All continuous variables were z-scored. *P*-values and 95% confidence intervals for the indirect effects were estimated using 10,000 bootstrap samples. Rows in bold indicate conditions where fixed effects are significant; \**P* < .05; \*\**P* < .01; \*\*\**P* < .001.

**Supplementary Table 6.** Indirect effect estimates of control input (GAD-7 total score) across time window parameters.

| Window size (sec) | Indirect effects of **speed** |  |  |  | Indirect effects of **angle** |  |  |  |
| --- | --- | --- | --- | --- | --- | --- | --- | --- |
|  | Beta coefficient | *P*-value | Lower bound of 95% CI | Upper bound of 95% CI | Beta coefficient | *P*-value | Lower bound of 95% CI | Upper bound of 95% CI |
| 3.0 | 0.139 | 0.238 | -0.044 | 0.395 | 0.009 | 0.862 | -0.035 | 0.140 |
| 3.5 | 0.135 | 0.265 | -0.063 | 0.388 | 0.018 | 0.662 | -0.022 | 0.156 |
| 4.0 | 0.129 | 0.264 | -0.062 | 0.368 | 0.009 | 0.844 | -0.056 | 0.131 |
| 4.5 | 0.138 | 0.239 | -0.052 | 0.380 | 0.004 | 0.930 | -0.065 | 0.110 |
| 5.0 | 0.132 | 0.248 | -0.043 | 0.383 | 0.008 | 0.898 | -0.066 | 0.130 |
| 5.5 | 0.147 | 0.204 | -0.022 | 0.389 | 0.016 | 0.828 | -0.085 | 0.164 |
| 6.0 | 0.167 | 0.169 | -0.018 | 0.397 | 0.006 | 0.925 | -0.102 | 0.130 |
| 6.5 | 0.153 | 0.192 | -0.023 | 0.388 | 0.002 | 0.965 | -0.123 | 0.117 |
| 7.0 | 0.148 | 0.197 | -0.029 | 0.379 | 0.000 | 0.943 | -0.122 | 0.091 |

*Note.* The mediation model used the GAD-7 total score as the input variable, the phase-averaged log-transformed latency measure as the output variable, and each dynamic feature as a mediator. Dummy-coded sex, age, and phase-averaged peak intensity of phase-congruent affect were controlled as covariates. All continuous variables were z-scored. *P*-values and 95% confidence intervals for the indirect effects were estimated using 10,000 bootstrap samples. All indirect effect estimates were not significant at $\alpha = .05$ .

**Supplementary Table 7.** Indirect effect estimates of control mediators (pure neural dynamics) across time window parameters.

| Window size (sec) | Indirect effects of **speed** |  |  |  | Indirect effects of **angle** |  |  |  |
| --- | --- | --- | --- | --- | --- | --- | --- | --- |
|  | Beta coefficient | *P*-value | Lower bound of 95% CI | Upper bound of 95% CI | Beta coefficient | *P*-value | Lower bound of 95% CI | Upper bound of 95% CI |
| 3.0 | 0.002 | 0.985 | -0.091 | 0.116 | 0.009 | 0.788 | -0.060 | 0.174 |
| 3.5 | 0.003 | 0.974 | -0.075 | 0.137 | 0.015 | 0.674 | -0.058 | 0.186 |
| 4.0 | 0.011 | 0.821 | -0.042 | 0.199 | 0.017 | 0.633 | -0.064 | 0.191 |
| 4.5 | 0.017 | 0.717 | -0.047 | 0.213 | 0.013 | 0.706 | -0.061 | 0.171 |
| 5.0 | 0.013 | 0.757 | -0.048 | 0.186 | 0.008 | 0.803 | -0.063 | 0.149 |
| 5.5 | 0.014 | 0.748 | -0.047 | 0.183 | 0.010 | 0.767 | -0.065 | 0.153 |
| 6.0 | 0.015 | 0.737 | -0.047 | 0.187 | 0.016 | 0.678 | -0.062 | 0.166 |
| 6.5 | 0.015 | 0.742 | -0.047 | 0.182 | 0.021 | 0.602 | -0.058 | 0.182 |
| 7.0 | 0.020 | 0.660 | -0.044 | 0.203 | 0.019 | 0.630 | -0.053 | 0.177 |

*Note.* The mediation model used the BDI-II total score as the input variable, the phase-averaged log-transformed latency measure as the output variable, and each dynamic feature calculated from the CEBRA-Time embeddings as a mediator. Dummy-coded sex, age, and phase-averaged peak intensity of phase-congruent affect were controlled as covariates. All continuous variables were z-scored. *P*-values and 95% confidence intervals for the indirect effects were estimated using 10,000 bootstrap samples. All indirect effect estimates were not significant at $\alpha = .05$ .

**Supplementary Table 8.**
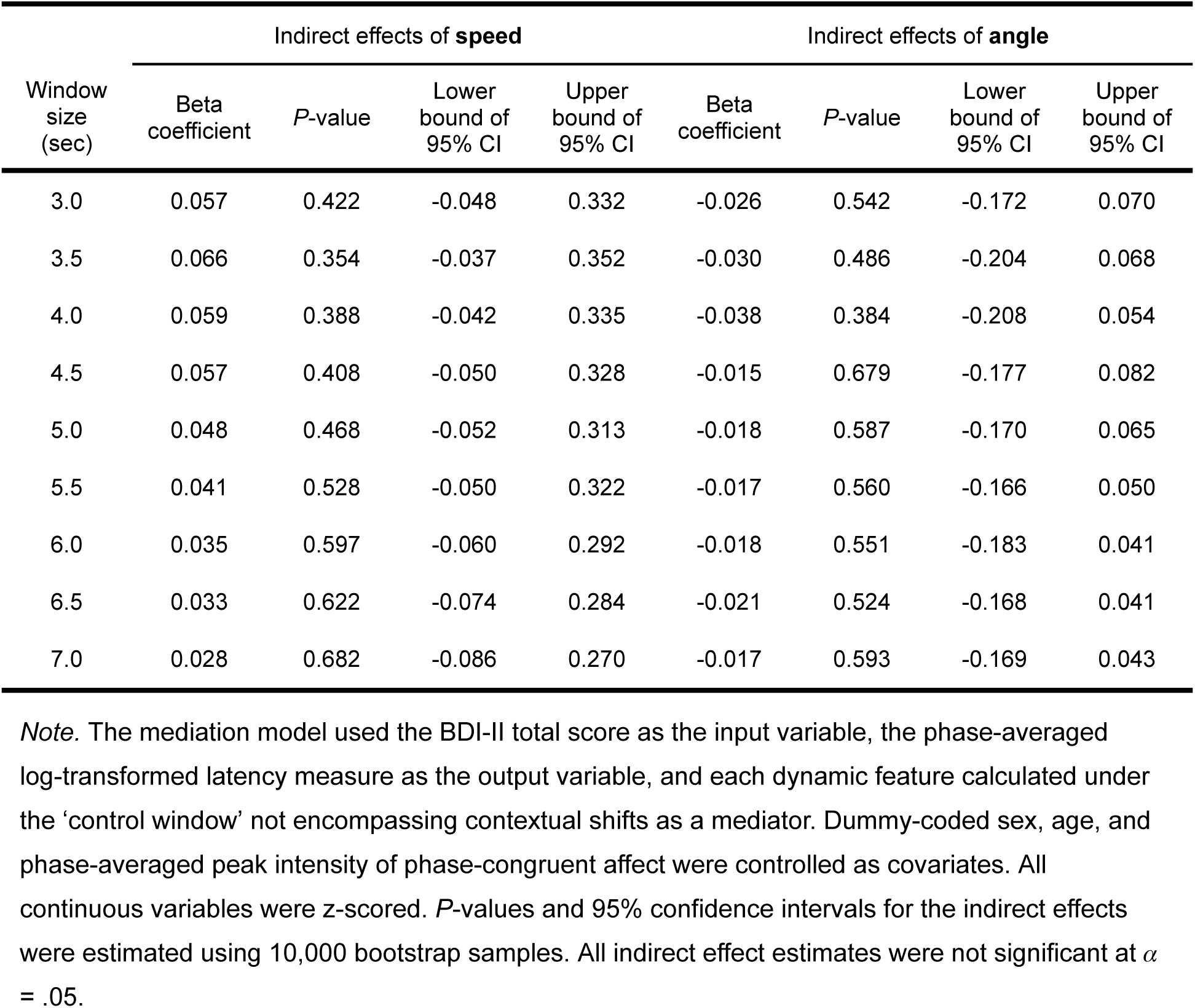
Indirect effect estimates under control time windows (no contextual shifts) across time window parameters.

